# Evaluating the likelihood for areas important for conservation to be recognized as Other Effective area-based Conservation Measures

**DOI:** 10.1101/2024.02.05.579011

**Authors:** Carly N Cook, Madhu Rao, Peter J Clyne, Vanessa Rathbone, Christian Barrientos, Antonio Boveda, Alex Diment, Jorge Parra, Valeria Falabella, Matthew Linkie, Deo Kujirakwinja, Stephane Ostrowski, Kirk Olson, Vardhan Patankar, Lovy Rasolofomanan, Hedley S Grantham

## Abstract

Other effective area-based conservation measures (OECMs) have expanded area-based conservation to recognize sites that deliver effective biodiversity outcomes even if not managed for conservation. Yet our ability to identify sites likely to qualify as OECMs remains limited. To address this gap, we established and tested a set of indicators to judge whether sites meet the essential criteria to be considered OECMs, evaluating a large, global sample of 173 important conservation areas: 81 potential OECMs and 92 nearby protected areas. We found that most potential OECMs were largely in good condition with the potential to achieve conservation outcomes, but none currently met all the OECM criteria. Formally designated protected areas in our dataset performed better but the majority also failed the criteria. With so many important conservation areas unable to deliver effective conservation outcomes, our findings raise important questions about how to ensure area-based conservation promotes positive and sustained outcomes for biodiversity.

## INTRODUCTION

There is consensus that halting the biodiversity extinction crisis and promoting the recovery of nature will require a significant expansion in conservation areas. Traditionally, area-based conservation has been focused on formal protected areas (PAs) as the principal tool to conserve biodiversity^1^. Internationally, there has been a significant increase in PAs since 2010^2^, promoted by targets set under the Convention on Biological Diversity (CBD)^3^. As the evidence for the scale of the twin biodiversity and climate crises become increasingly clear^4^, there is growing momentum for bolder area-based conservation targets^5^.

In response to calls for more ambitious protection targets, in 2022 the CBD Global Biodiversity Framework (GBF) committed signatory countries to protect at least 30% of land and sea by 2030^6^. An analysis of progress towards Aichi Target 11 revealed that by 2020, the world not only fell short of the 17% protection target for land and 10% for the sea, but also failed to meet the qualitative elements of Target 11: “ecologically representative, well-connected, effectively and equitable managed areas”^2^. There is now a monumental challenge before the global community to meet all aspects of Target 3 within the GBF by 2030. This will require both improving the effectiveness of existing PA networks and rapidly expanding protection for biodiversity in areas that deliver successful conservation outcomes beyond PAs.

In 2010, ‘other effective area-based conservation measures’ (OECMs) were introduced as a new form of area-based conservation (i.e., CoP14, Decision 7;^3^). However, it was not until 2018 that OECMs were defined, characterized as areas “governed and managed in ways that achieve positive and sustained long-term outcomes for the in-situ conservation of biodiversity…”^7^. An important feature that distinguishes OECMs from PAs is that biodiversity conservation need not be the primary objective but can instead be a secondary objective or indirect benefit (ancillary objective) of other objectives^8^. The other major departure from PAs is the requirement that OECMs provide effective conservation outcomes^7^. In contrast, PAs are defined by their management objectives rather than explicit biodiversity outcomes, and not all PAs are effective at halting biodiversity loss^9,10,11^. Given this important conceptual shift, the delay in providing a definition and criteria for OECMs^7^ and associated guidance^8,12^, along with the complexity involved in interpreting and applying the criteria across diverse contexts^13^, have created challenges for identifying OECMs. To date 870 OECMs have been formally registered across nine jurisdictions^14^ and it is unclear whether some of these areas genuinely meet the relevant criteria (e.g., marine refuges in Canada^15^).

There remains uncertainty about how to translate the broad criteria and general guidance for OECMs into specific recommendations for which areas should be counted towards area protection targets^16^. A site-level assessment tool^12^ has been developed to complement the CBD definition and existing guidance on recognizing and reporting OECMs^8^, yet considerable doubt remains about what is required to demonstrate that specific areas meet the necessary criteria^13,17,18^. Identifying high performing areas needs detailed guidance, formulated from an evidence-base developed on the back of diverse case studies identifying candidate sites, and monitoring and reporting on recognized OECMs^19^. Likewise, collaboration is needed across a wide range of stakeholders to ensure the evidence-base and policy mechanisms are in place at national and sub-national levels to recognize areas governed by both state and non-state actors^21^. This evidence could help maximize the benefits of area-based strategies^22^.

Given the expectation that OECMs should achieve positive and sustained conservation of biodiversity, a focus on OECMs could renew the emphasis on the effectiveness of area-based conservation more generally^18,23^, rather than just the total area protected^22,24^. There are many actors whose management practices support positive conservation outcomes^16,25^, including Indigenous Peoples and local communities^13,26^ and private landholders^27^. Applied within a systematic conservation planning framework, OECMs have the potential to expand the representation of biodiversity and promote connectivity among existing PAs^13^. Although the limited engagement with the evaluation of possible and existing OECMs means that realizing these potential benefits remains uncertain^17,19,28^.

To avoid the potential pitfalls, and realize the promised benefits of OECMs, it is essential that the emphasis is firmly on effective conservation outcomes and developing a pathway to support sites to move toward this goal^20^. It has been stressed that identifying OECMs requires a site-by-site approach to assessing individual areas against the CBD criteria^8,12^, but only a handful of studies have attempted to use the IUCN guidance to screen possible OECMs^17^, and the methods used to determine whether sites meet the definition rely largely on expert judgment (e.g.,^25^). Therefore, more than a decade after they were first conceived, critical gap in the conservation science required to screen possible OECMs remain.

To help fill this gap, we assessed a large global sample of areas important for conservation against the essential aspect of the OECM definition - achieving “positive and sustained long-term outcomes for the in-situ conservation of biodiversity”. Importantly, we sought to assess the performance of potential OECMs and formal PAs in the same regions. As conservation areas within an existing system and associated with a well-established framework (i.e.,^29^), PAs offer a valuable point of comparison for the potential OECMs. We provide the most comprehensive, global assessment of important conservation areas using the criteria for OECMs derived from the CBD definition, interpreted according to the guidance for recognizing OECMs^8^ and the associated site-level assessment tool^12^. Our findings provide critical insights into how conservation areas perform when the emphasis is placed on sustained biodiversity outcomes.

## METHODS

### Indicator development

To evaluate sites, we drew on the formal definition of OECMs to demonstrate *positive and sustained long-term outcomes for the in-situ conservation of biodiversity*^7^. Drawing on the guidance from the IUCN, we developed a number of indicators that align with the descriptions used to complement this definition by interpreting and expanding on the criteria^8,12^. The first two criteria in the site-level assessment tool identify potential OECMs: Criterion 1 – not a recognized PA and Criterion 2 – “*likely* to support important biodiversity”. We used these criteria to identify possible sites (see Site Selection). The indicators developed for each of the remaining criteria were designed to translate the relevant guidance (i.e., the criteria and their associated ‘elements’) into statements with categorical responses. While the site-level assessment tool^12^ was not available at the time our indicators were developed, the indicators within our questionnaire align well with the criteria and guidance in the site-level assessment tool (Table 1).

**Table 1:**
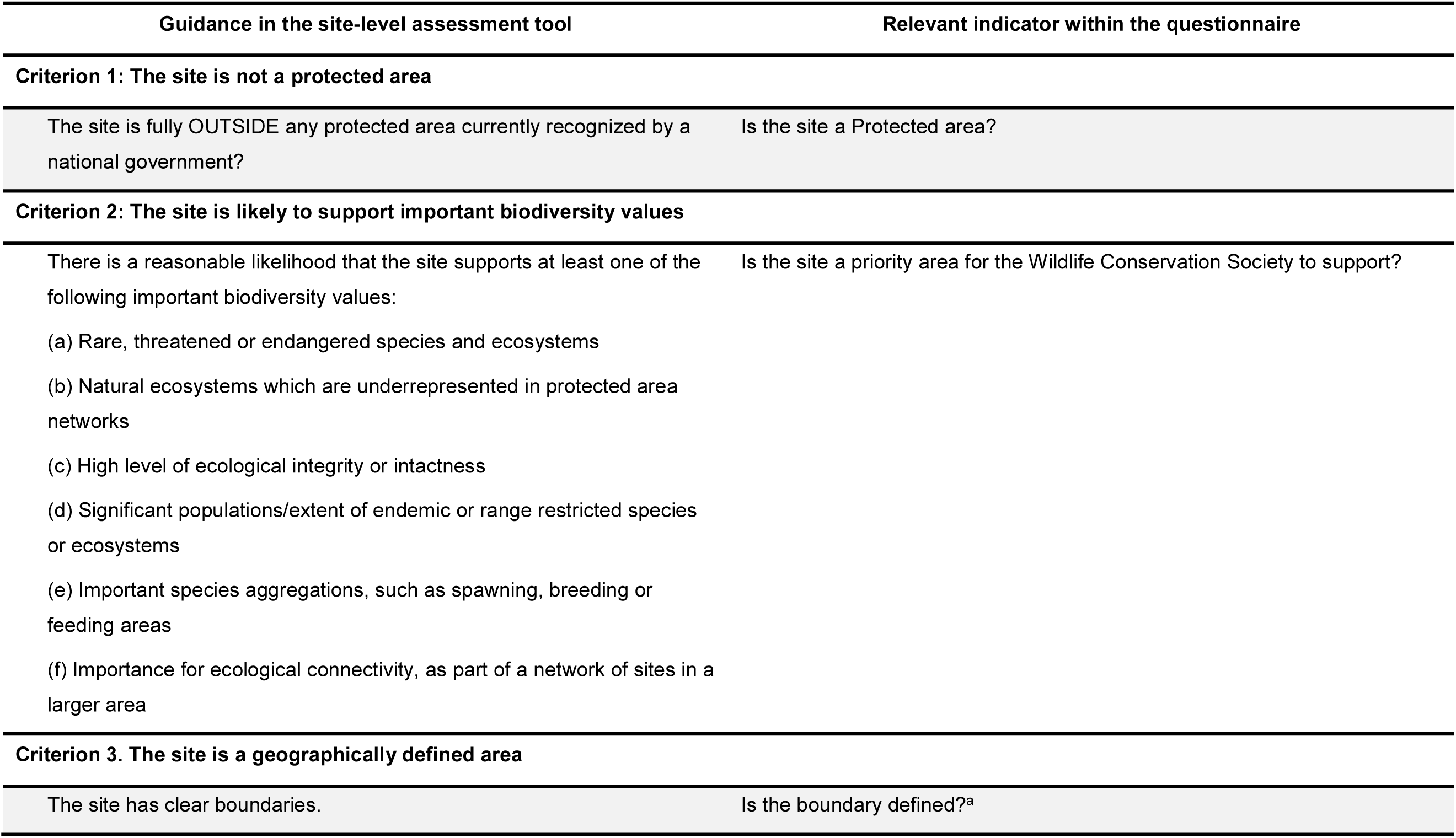
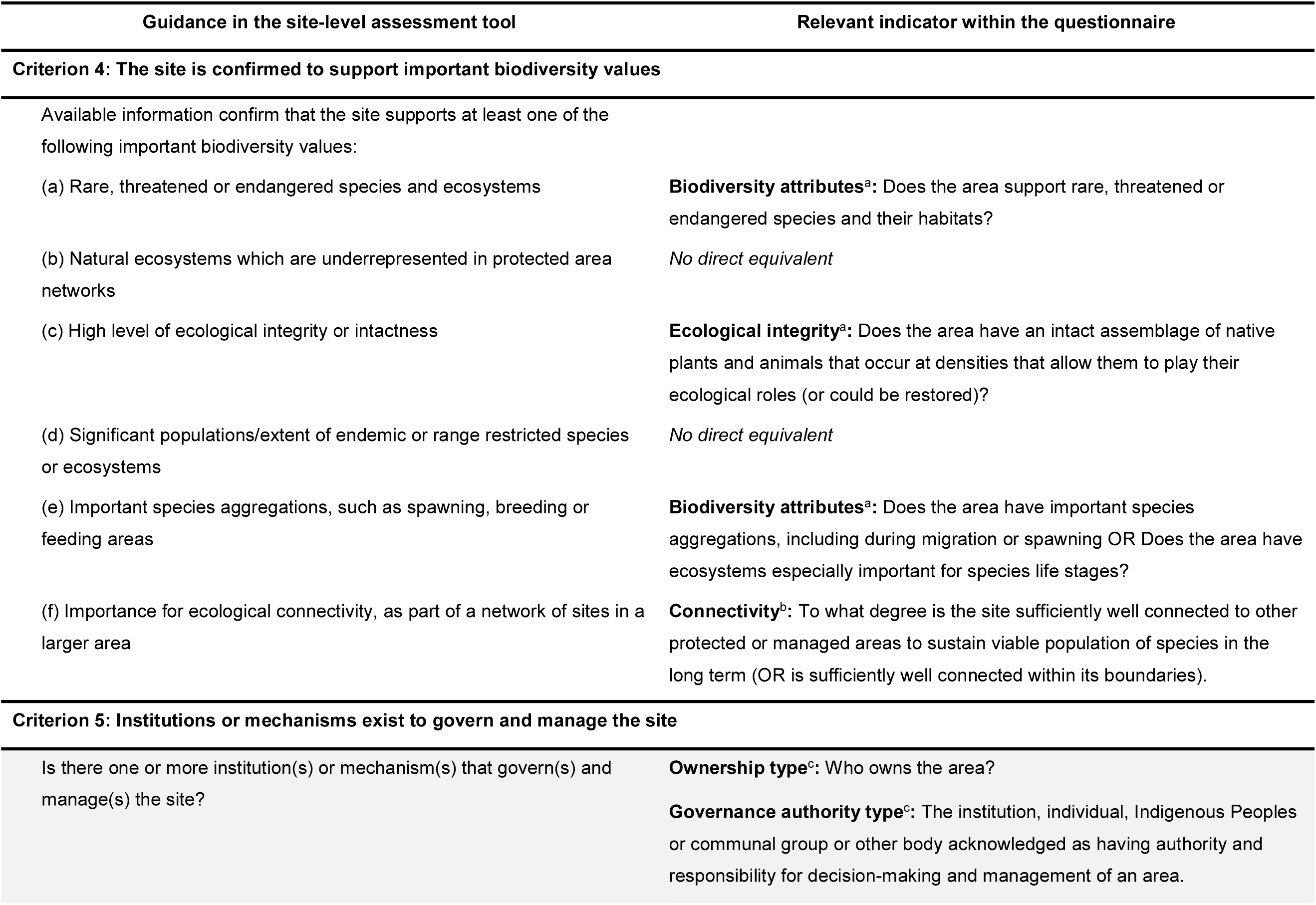
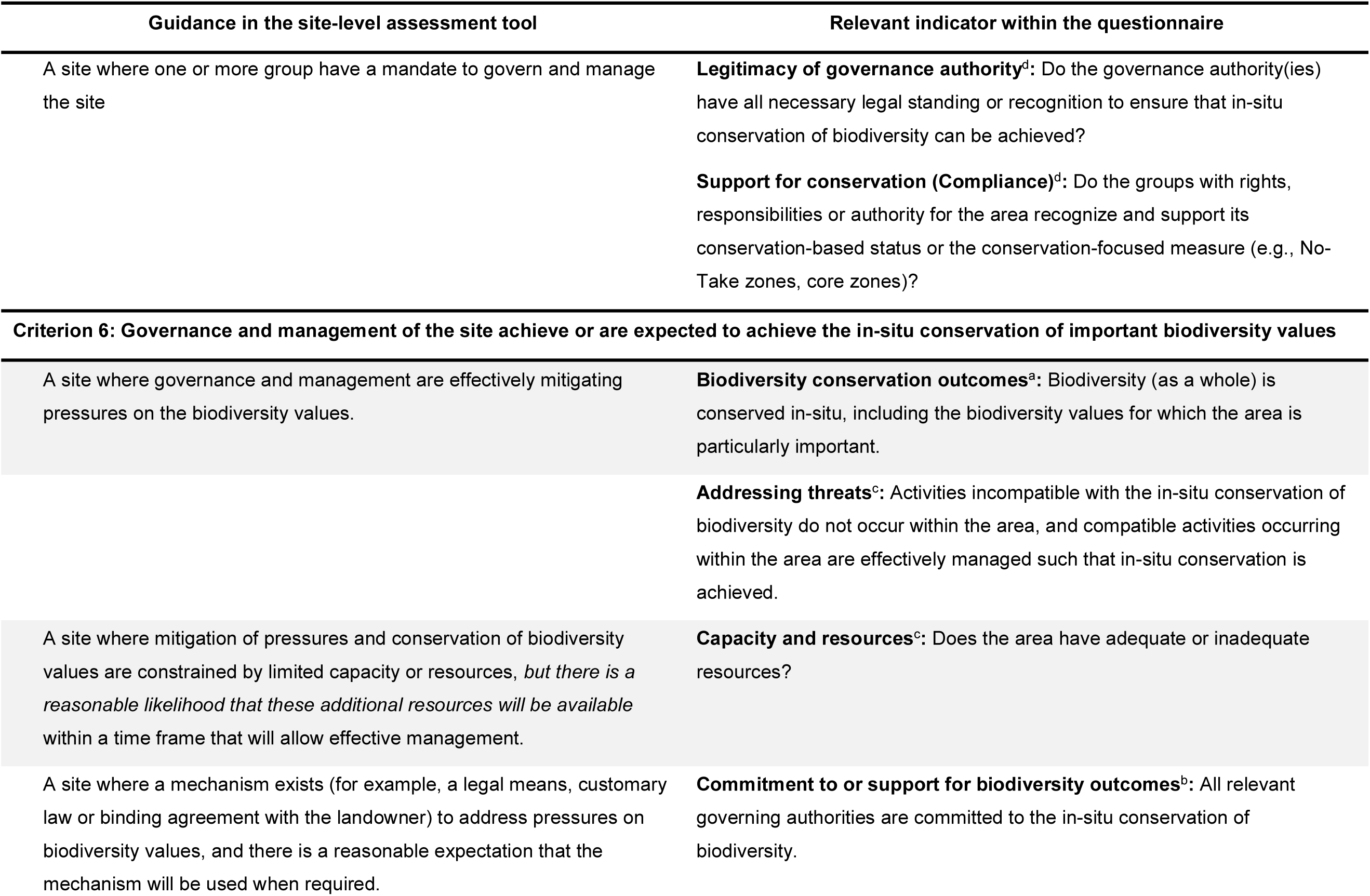
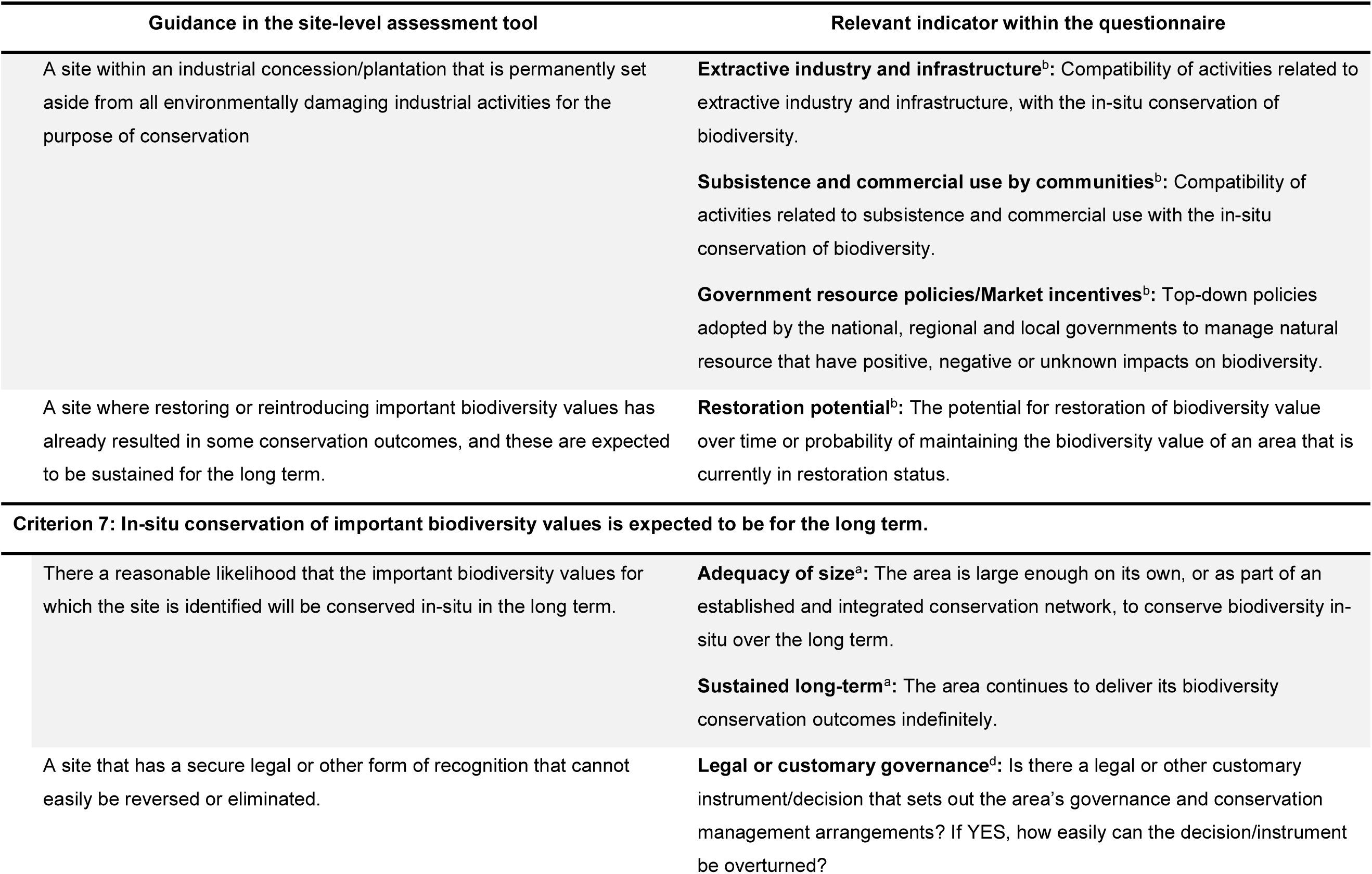
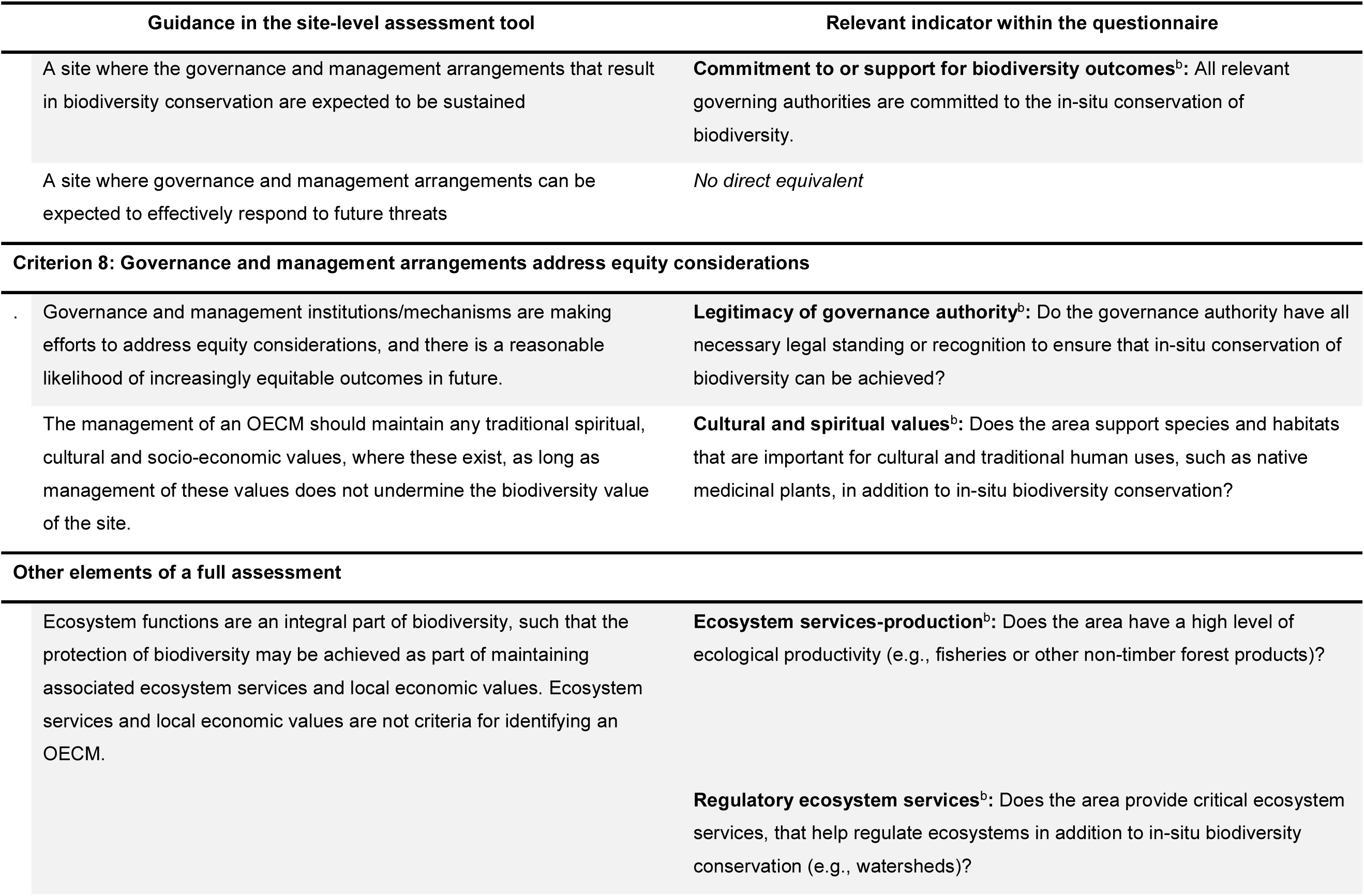
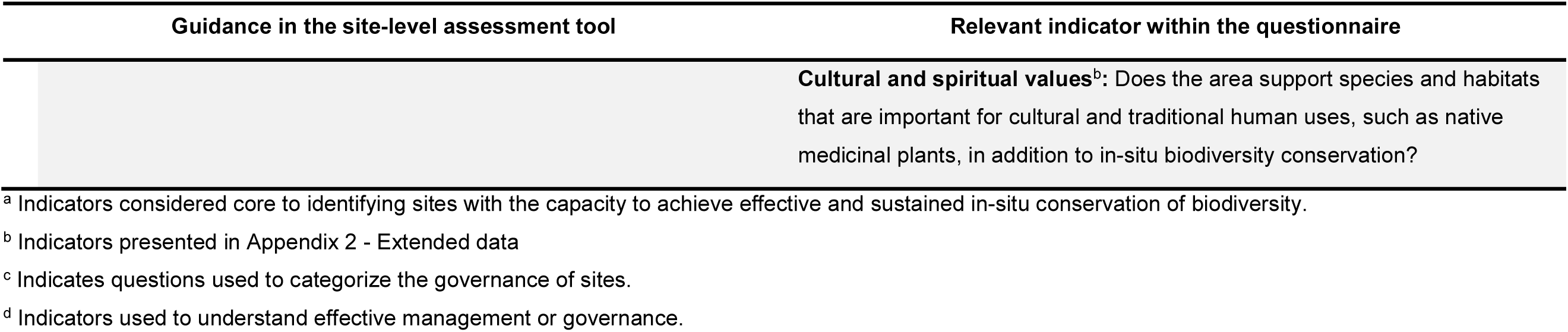
The indicators used to evaluate potential other effective area-based conservation measures (OECMs) and how they relate to the criteria and associated guidance within the site-level assessment tool^10^. Details of each indicator are provided in Appendix S1. Criteria 1 and 2 were used to identify potential OECMs for data collection.

We developed 27 indicators that span all eight criteria in the site-level assessment tool (^12^; Table 1). Each indicator was expressed as a statement that summarized what is required to meet the criterion or relevant explanation within the IUCN guidance (Table 1). The range of possible responses were designed to capture conditions where sites fully, partially or did not meet the criterion (Appendix S1), consistent with the site-level assessment tool.

The indicators were used to create a questionnaire, including descriptive information about the site. This included the area (km^2^), whether the boundary of the site defined (i.e., Criterion 3: geographically defined area; Table 1), the relevant biome and dominant ecosystems present, and the type of site (e.g., community forest, territorial lands).

Our primary focus was to assess key aspects of Criterion 4: the presence of important biodiversity; Criteria 6: effective *in-situ* conservation of biodiversity; and Criterion 7: the ability to sustain conservation outcomes long-term. Therefore, we focus on three key indicators relating to in-situ conservation of important biodiversity and three indicators focused on long-term conservation outcomes, supplemented by the remaining indicators focused on evaluating the management and governance of the site (Table 1). Questions about the ownership (who owns the site) and governance of the site (who has the authority and responsibility for decision-making and management) (Criterion 5; Table 1) were asked to classify the sites in terms of the IUCN governance types^28^.

### Site selection

All sites were drawn from regions that the Wildlife Conservation Society (WCS), an international conservation NGO, had identified as priorities for nature conservation. These areas are ecologically representative of a range of global ecoregions and contribute to conservation in at least two of the following ways: containing disproportionately high levels of important biodiversity; maintaining intact ecosystems; protecting important areas for iconic or charismatic megafauna; providing refugia for unique wildlife; and making important contributions to climate change resilience^39^.

These regions are almost always a mosaic of formal PAs and other areas important for biodiversity conservation. Within these mosaics, we selected places that meet the definition of “potential OECMs” based on the site-level assessment tool (i.e., satisfy Criterion 1 and Criterion 2)^12^. The decision to target both PAs and potential OECMs was made to reflect the landscape-scale approach WCS applies to their conservation efforts, offering a unique opportunity to contrast the current effectiveness of conservation areas with and without formal legal protection.

Individual sites in the sample were selected by local experts, who were field-based conservation scientists and practitioners with deep knowledge of the ecological, social and economic contexts of the selected sites. Experts were asked to identify at least one area that could be a potential OECM and one PA. When selecting potential OECMs, experts could choose sites that function to conserve biodiversity in their own right, or by supporting conservation efforts in nearby PAs (i.e., enlarging, buffering or connecting existing PAs). Priority was given to selecting sites for which experts had the most detailed knowledge.

### Data collection

The questionnaire was used to evaluate 173 conservation areas across 23 countries (Figure 1). The sites were evaluated by 18 local experts, who nominated relevant PAs and potential OECMs for which they had detailed knowledge, as per the site-level assessment tool guidelines^12^. A separate questionnaire was completed for each site. Given the nature of WCS country programs, some experts were able to provide responses for multiple sites where they are actively involved in supporting governance and management.

**Figure 1.**
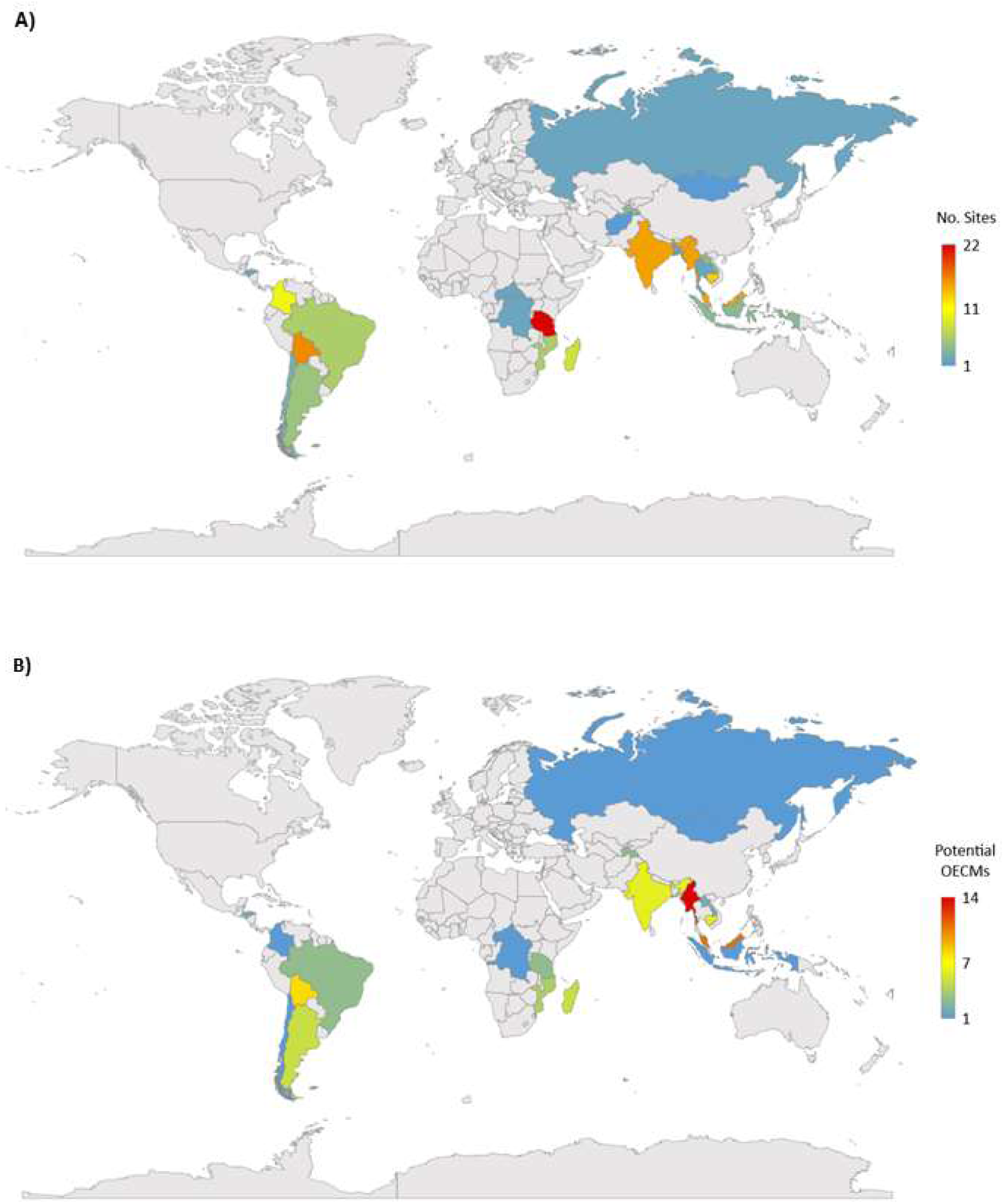
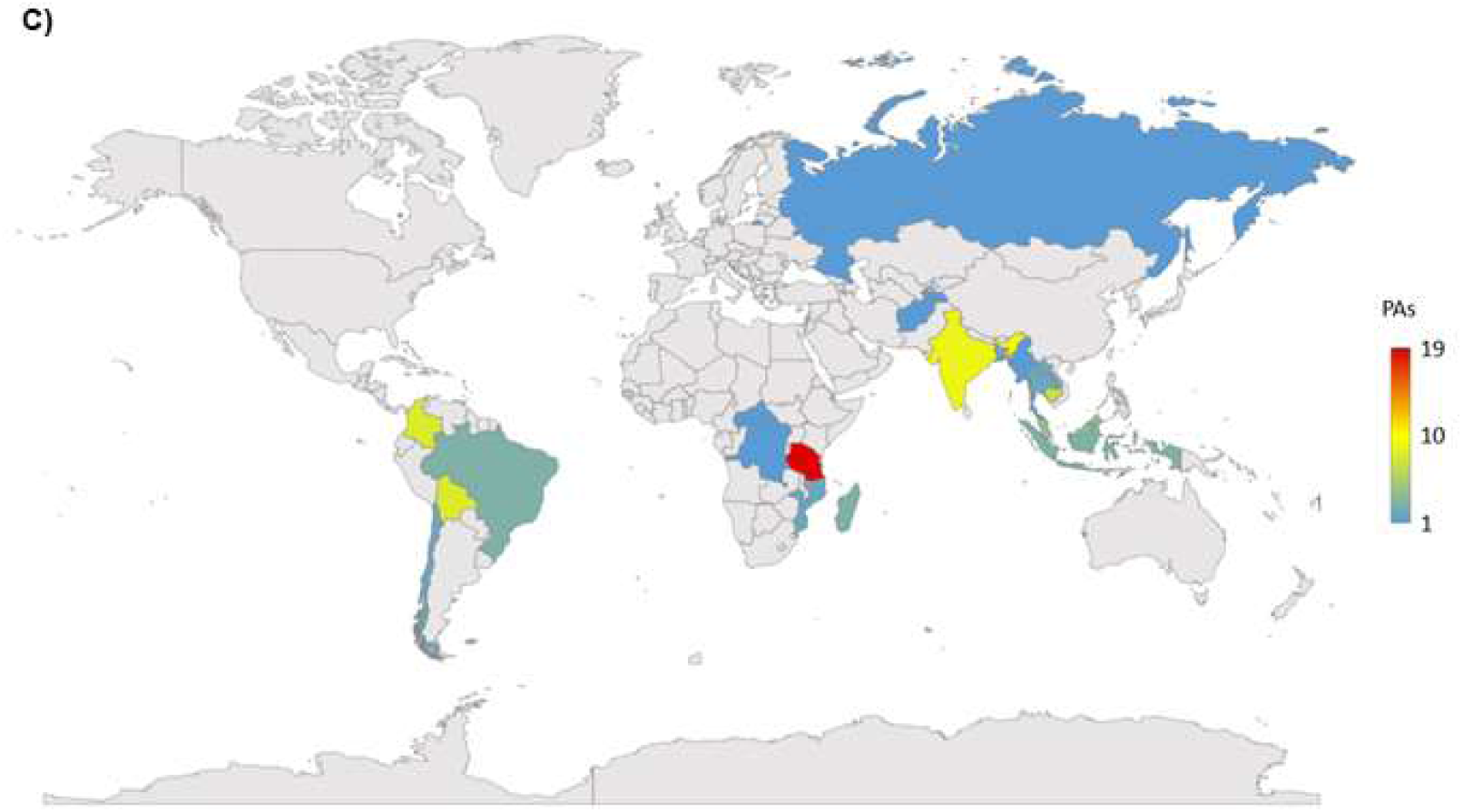
The geographic distribution of sites in the data set. Panel (A) shows the total number of sites per country (n=173). Panel (B) shows the number of potential OECMs per country (n=81). Panel (C) shows the number of protected areas (PAs) per country (n=92).

Each expert was contacted and asked if they were willing to participate. If they agreed, they were provided with a questionnaire and detailed instructions they could use to complete the questionnaire independently. Experts could also opt to be supported to complete the questionnaire by a member of the research team. In these cases, the questionnaire functioned more like an interview, where experts were asked which response they felt best reflected the site. In addition to acting as local experts, respondents could draw on other verified sources to support their responses for each indicator, including scientific studies, local laws and regulations, and relevant traditional and Indigenous knowledge or that of other local experts^12^. Questionnaires were completed between August 2021 and August 2022. Experts from four countries opted not to provide data on potential sites due to a lack of sufficiently detailed knowledge.

### Data analysis

To examine whether potential OECMs performed differently to PAs across the different indicators we used Mann-Whitney U non-parametric tests^40^. Each indicator was converted to an ordinal response variable, scaled between 1 (lowest performance) and 3 (highest performance) (Appendix S1). Mann-Whitney U non-parametric tests were also used to compare the performance of marine and terrestrial sites across the indicators. All statistical analyses were conducted in the software program SPSS Version 27. P-values <0.05 were considered to indicate statistical significance.

## RESULTS

Evaluations were completed for 173 sites (92 PAs and 81 potential OECMs) across 23 countries (Figure 1), including important marine and terrestrial areas (Extended data; Figure S1). While a third of sites were in the marine realm (n=55), the proportion of potential OECMs versus PAs evaluated was consistent across realms (44% and 48% of marine and terrestrial sites respectively).

The sample represented sites with diverse ownership types and governance arrangements (Figure 2). The majority of sites were owned by state actors (70%), with communal (13%), private (2%) and multiple ownership (15%) arrangements being less common. While the ownership of potential OECMs and PAs was similar, communal governance arrangements were more common for potential OECMs (Figure 2A). Governments were responsible for the management of half the sites (51%), and 28% of sites had shared governance arrangements. Despite only representing a fifth of the sample (19%), most sites managed by Indigenous and local communities were potential OECMs (Figure 2B).

**Figure 2.**
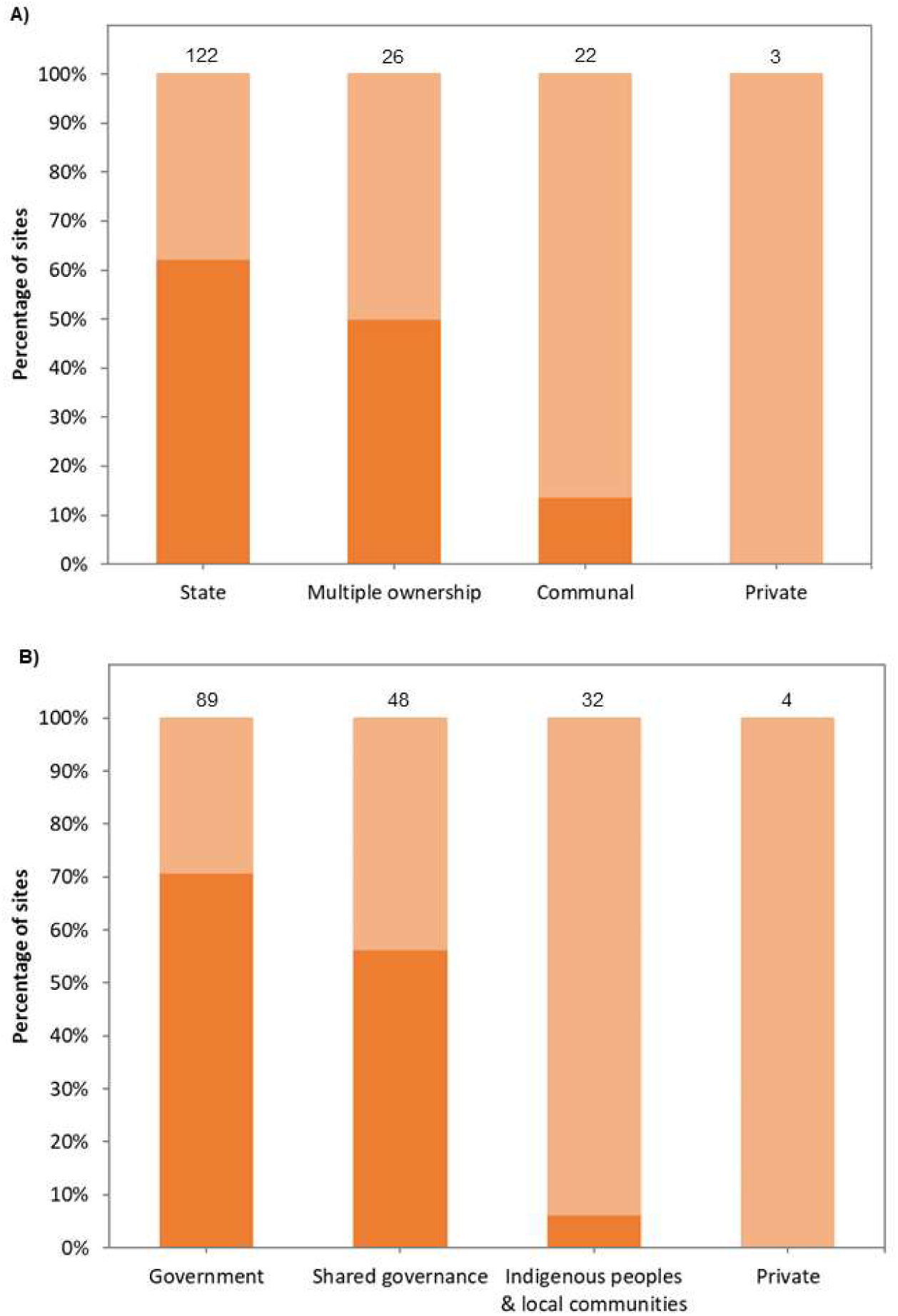
The governance arrangement for all sites in the dataset according to (A) who owns the relevant land or waters; and (B) the governance authority who has the responsibility for making decisions about the management of an area. Numbers denote the number of sites. Dark bars = protected areas. Light bars = potential OECMs.

By far the most common governance arrangements for PAs were state-owned sites managed by government actors (Figure 2). Potential OECMs had a much more diverse set of governance arrangements, with approximately a third of sites state-owned with management by government actors (30%) (Figure 2). Where potential OECMs were managed by Indigenous and local communities, almost a quarter were communally-owned (22%) and a sixth were state-owned (14%) areas (Figure 2).

The types of sites and extent to which biodiversity conservation was an objective for the management of potential OECMs varied widely (Figure 3). Most sites had biodiversity conservation as a secondary management objective (e.g., wildlife areas, forestry concessions, community-managed forests or fisheries areas). Almost one-third of sites had biodiversity as the primary objective (e.g., nature reserve or Biosphere reserve) and three sites had biodiversity conservation as an ancillary outcome (e.g., military lands) (Figure 3).

**Figure 3.**
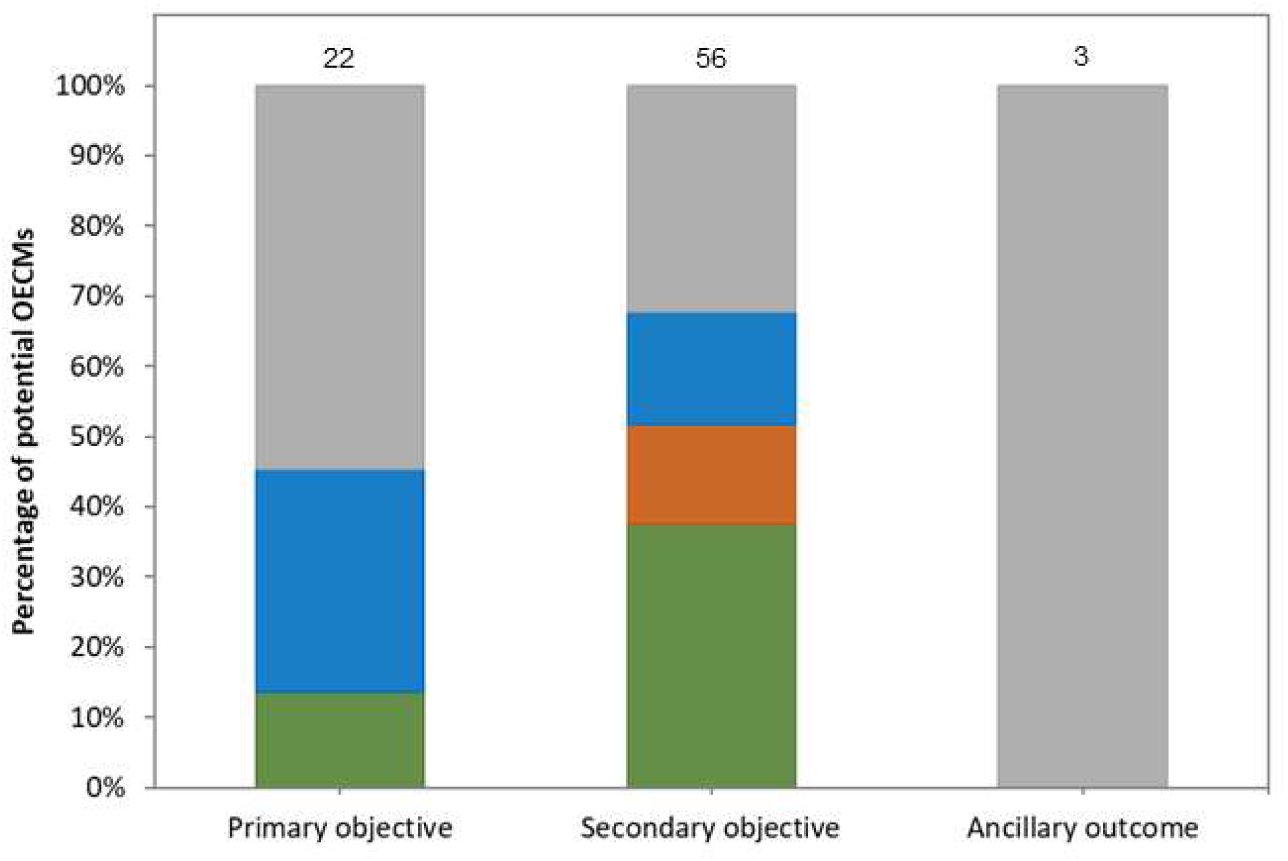
Distribution of where conservation management objectives rank within the potential Other Effective area-based Conservation Measures in our sample, and the broad types of areas included. Green = forest management areas. Brown = wildlife or game management areas. Blue = fisheries management or marine conservation areas. Grey = other.

### Essential Characteristics of Other Effective Area-based Conservation Measures

#### CBD Criterion 3: A geographically defined area

Experts reported that 70% (n=57) of potential OECMs had defined boundaries, relative to 89% (n=82) of PAs. As such, 19% of the sample failed the first step in a full assessment. Neverthless, we analyzed all of the sites, independent of whether or not they had clear boundaries.

#### CBD Criterion 4: Important biodiversity values and associated ecosystem services

Experts confirmed that all sites were able to support at least one of the important biodiversity values set out in Criterion 4. In our sample, 91% of potential OECMs and 97% of PAs were reported to support rare or threatened species and their habitats, important species aggregations, and/or important ecosystems (Criteria 4a & 4e, Table 1). The majority of sites had high levels of connectivity (59% of potential OECMs, 65% of PAs; Figure S2) being considered sufficiently well-connected to other protected or managed areas to sustain viable population of species (Criterion 4f, Table 1). The small number of sites that were not reported to be important for specific species and their habitats were forested areas which reported their values as species richness and ecosystem services.

PAs had significantly higher levels of ecological intactness than potential OECMs (*U*=4217.00; z=2.10; p=0.035), with almost a third (28%) of potential OECMs deemed to have native biota at densities below what would be required to allow them to play their ecological roles (Criterion 4c, Table 1; Figure 4A), Marine sites were more likely to have intact assemblages than terrestrial sites (*U*=3915.50; z=3.08; p=0.002).

**Figure 4.**
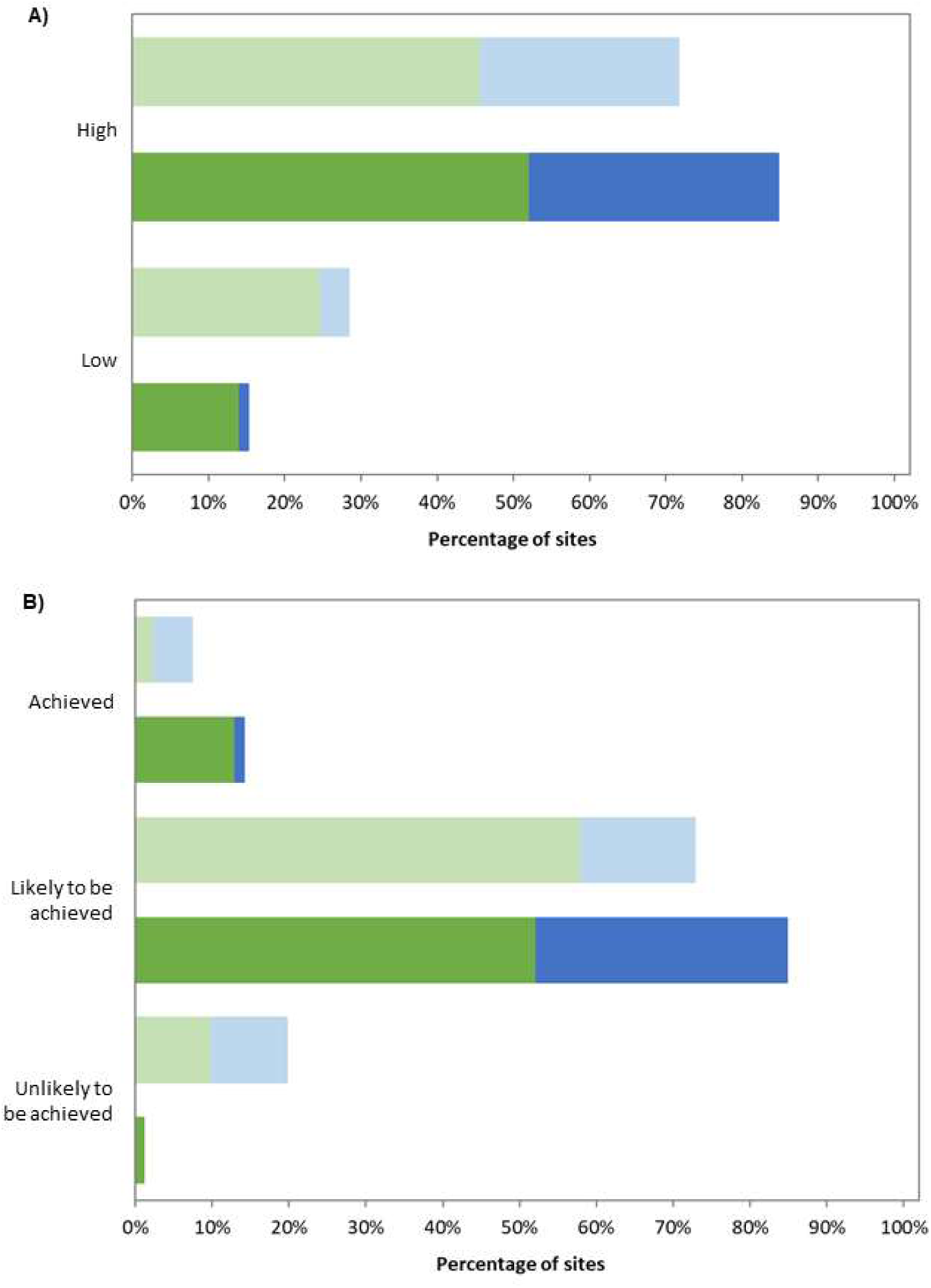
The percentage of sites where (A) the integrity of species assemblages is high, and (B) in-situ conservation of biodiversity has been achieved. Light bars = potential OECMs (n=81); Dark bars = PAs (n=92). Green = terrestrial sites (n=118); Blue = marine sites (n=55).

While not considered a key criterion for OECM status, the vast majority of sites (>80%) were reported to be important for both regulatory and production ecosystem services. Most sites were also important for cultural and spiritual services (77% of potential OECMs; 66% of PAs), although the status of cultural ecosystem services was unknown for many sites (19% of potential OECMs; 24% of PAs).

#### CBD Criterion 5: Institutions or mechanisms exist to govern and manage the site

The sites spanned the full range of governance types recognized by the IUCN (Figure 2^29^). The site level assessment tool states that the group(s) responsible for the site must have a mandate to govern and manage the site. We found that 66% of potential OECMs had a governing authority with the necessary legal standing or recognition to ensure the conservation of biodiversity, significantly lower than the 98% of PAs (*U*=4661.00; z=5.49; p<0.001; Figure S3). This was a particular problem for potential marine OECMs, where only 35% of sites had recognized governing authorities, significantly lower than those in terrestrial environments (*U*=329.00; z=-3.41; p=0.001).

The majority of sites had groups that recognize and support its conservation status (Figure S3). However, 21% of potential OECMs had at least one governing authority who did not support in-situ conservation of biodiversity (Figure S3).

#### CBD Criterion 6: Governance and management of the site achieve or are expected to achieve the in-situ conservation of important biodiversity values

##### Effective in-situ conservation of biodiversity

While 73% of potential OECMs were considered likely to achieve their conservation outcomes at some point in the future, only 7% were considered to have already achieved effective in-situ conservation of the biodiversity for which the site was important (Figure 4B). PAs were considered significantly more likely to achieve effective conservation outcomes (*U*=3914.00; z=3.98; p<0.001); nevertheless, only 14% were considered to have already achieved effective in-situ conservation of biodiversity (Figure 4B).

##### Preventing and mitigating threats to biodiversity

Several indicators addressed whether the governance and management of a site prevent and mitigate threats to the site’s important biodiversity values (Figures S4-S5). Threats were reported to have been identified and were being effectively addressed for only 9% of potential OECMs and 17% of PAs, but were being at least partially addressed in the majority of areas (Figure 5A). However, significantly more potential OECMs were considered to have threats negatively impacting biodiversity (Figure 5A; *U*=3605.00; z=2.15; p=0.032). Unsurprisingly, only a fraction of PAs or potential OECMs were considered to have sufficient resources (i.e., staff and operational budget) to sustain long-term conservation outcomes (Figure 5B; *U*=3869.00; z=1.35; p=0.117).

**Figure 5.**
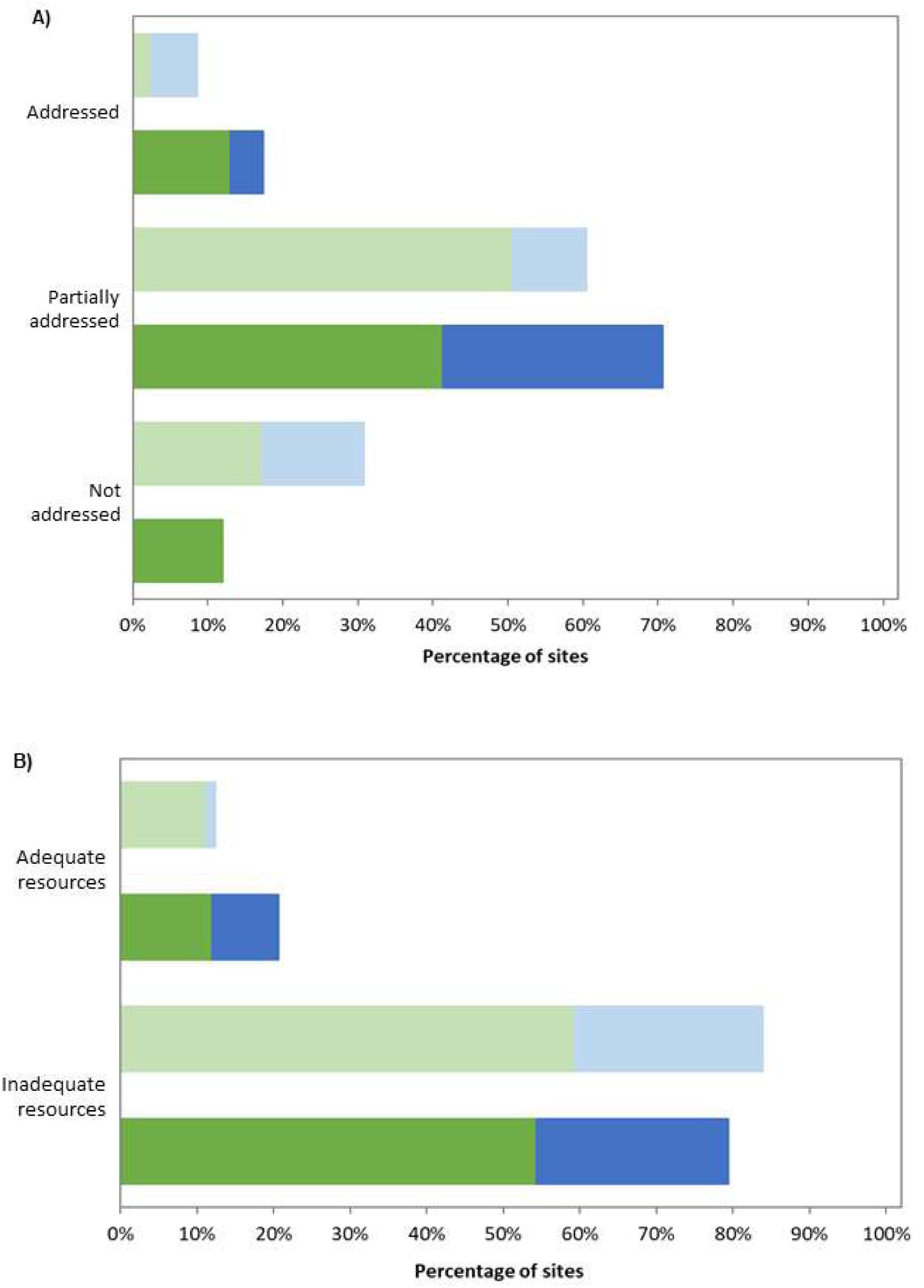
The percentage of sites where (A) threats are identified and effectively addressed, and (B) sites have adequate resources to sustain long-term conservation outcomes. Light bars = potential OECMs (n=81); Dark bars = PAs (n=92). Green = terrestrial sites (n=118); Blue = marine sites (n=55).

#### CBD Criterion 7: In-situ conservation of important biodiversity values is expected to be sustained long term

##### Long-term conservation outcomes

While biodiversity outcomes are a critical pillar of OECMs, the governance and management structures in place also need to be able to support long-term, sustained outcomes^7^. Potential OECMs were less likely to be large enough to achieve the in-situ conservation of biodiversity over the long-term than PAs (Figure S6; *U*=4620.50; z=3.65; p<0.001). Most PAs were deemed to be large enough, either alone (52%) or in combination with nearby areas (38%) (Figure S6). The vast majority of potential OECMs were considered only to be viable in combination with nearby areas (78%), but 21% were considered large enough to achieve the long-term conservation of important biodiversity in their own right (Figure S6).

Approximately half of potential OECMs and PAs were believed to be likely to achieve sustained biodiversity outcomes; overall significantly more PAs were expected to achieve long-term outcomes (*U*=3801.00; z=3.84; p<0.001; Figure 6A).

**Figure 6.**
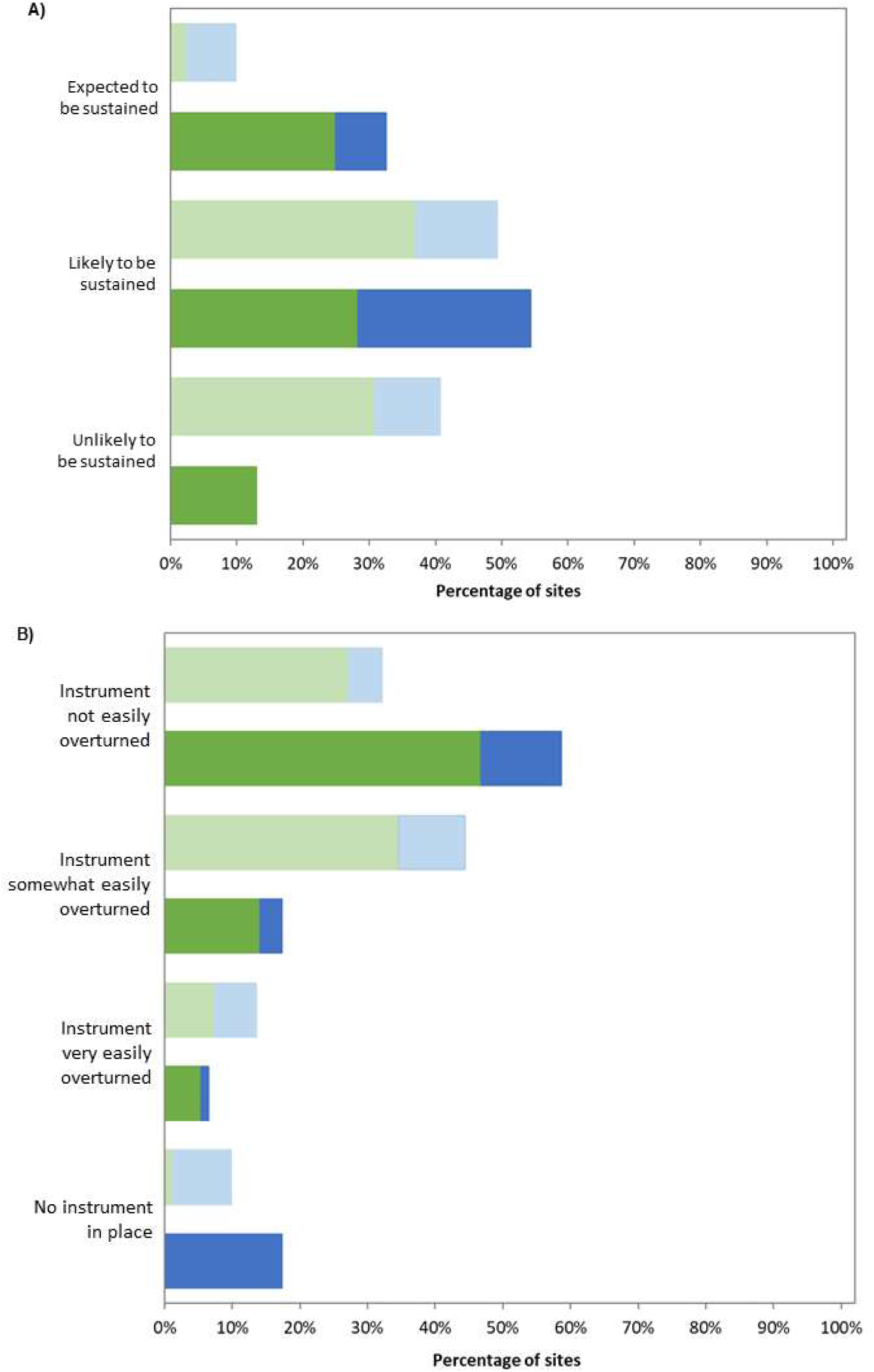
The percentage of sites where (A) biodiversity conservation outcomes are likely to be sustained long-term, and (B) there is a secure instrument in place to support governance and management. Light bars = potential OECMs (n=81); Dark bars = PAs (n=92). Green = terrestrial sites (n=118); Blue = marine sites (n=55).

Potential OECMs were significantly less likely than PAs to have monitoring in place to track conservation outcomes (*U*=2272.00; z=-2.26; p=0.024), with less than a third of sites monitoring the important biodiversity that occur there (Figure S7).

Concerningly, approximately 69% of potential OECMs had insecure governance and conservation management arrangements, either because there was no instrument in place or the instrument in place could be easily or somewhat easily overturned (Figure 6B). Conversely, while most PAs had secure arrangements in place, one third were at least somewhat insecure.

##### Overall capacity to achieve effective and sustained in-situ conservation of biodiversity

The site-level assessment tool states that a site must meet all of the criteria to be a “confirmed OECM”, while those that only partially meet any criteria remain a “candidate OECM”^12^. When considering the performance of sites across the essential indicators relating to achieving effective and sustained in-situ conservation, we found that none of the 81 potential OECMs in our dataset could be deemed OECMs (Figure 7A). However, including sites that could be considered candidate OECMs (i.e., sites that scored in the intermediate range for some indicators), this number rose to 29 sites (Figure 7B). While only four of the PAs in our sample were deemed to meet the essential indicators to be OECMs, two-thirds (n=61) could be considered equivalent to candidate OECMs (Figure S8).

**Figure 7.**
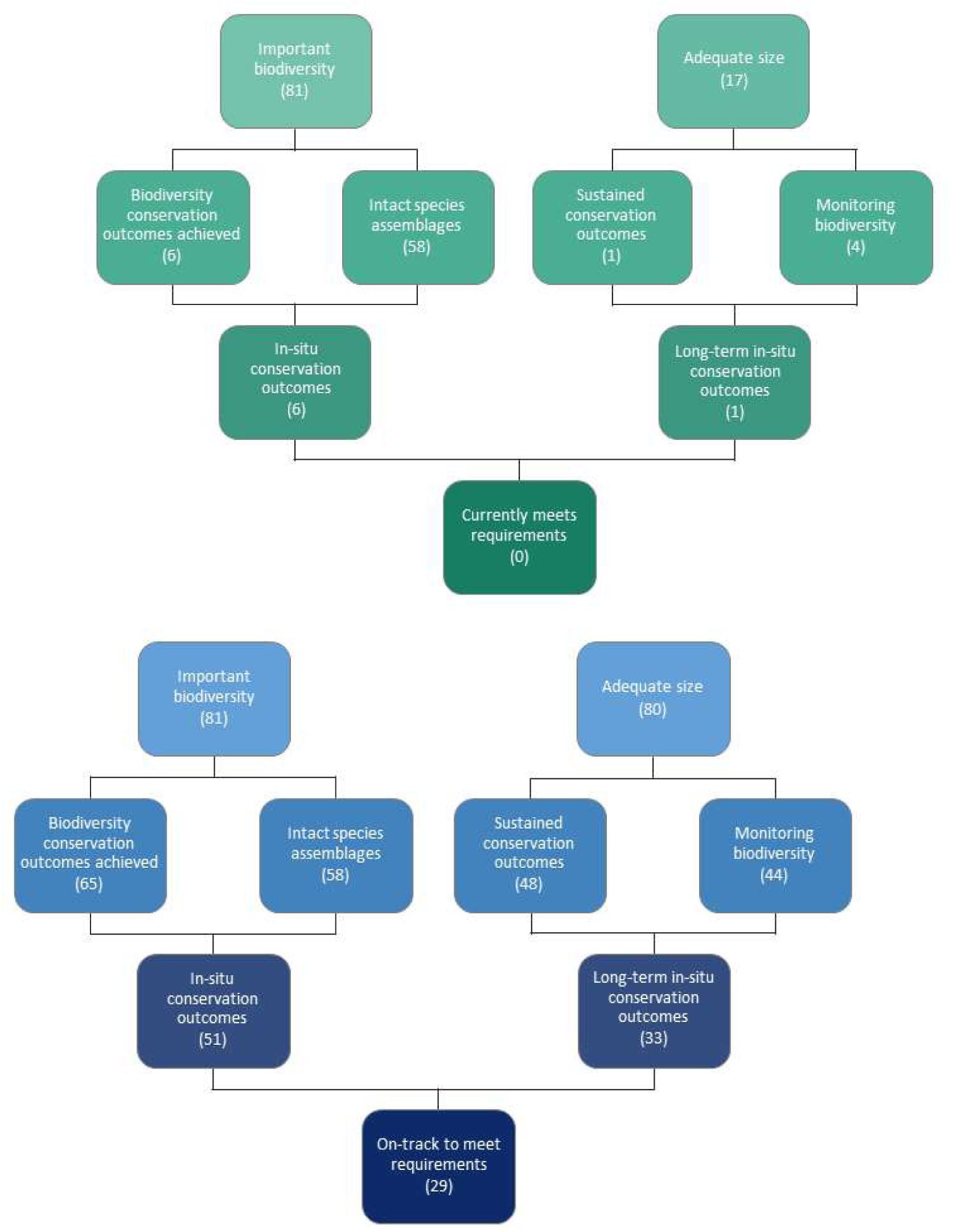
The number of potential OECMs that (A) currently or (B) might be on track to achieve effective and sustained in-situ conservation of biodiversity. Sites in Panel (A) could be equivalent to OECMs and sites in Panel (B) could be considered candidate OECMs. Numbers denote the number of sites, with 81 potential OECMs in total.

## DISCUSSION

We set out to evaluate a wide range of conservation areas against the criteria for OECMs. Our data represents a large and diverse set of important conservation areas from around the world, including both formal PAs and potential OECMs, comprising multiple governance types across marine and terrestrial environments. As the most significant effort to screen potential OECMs against the formal criteria, these data provide important insights into the challenges associated with implementing OECMs in practice.

### Demonstrating areas meet the required OECM criteria

Concerns have been expressed that the definition of OECMs sets a very high standard for sites^13^. We found that almost all conservation areas in our sample failed to meet the criteria of achieving sustained outcomes for the in-situ conservation of biodiversity (Figure 7), including the PAs (Figure S9). However, many of these conservation areas were good candidate OECMs because, while they were not yet achieving in-situ conservation outcomes, they were on-track to achieve positive conservation outcomes in the future (Figure 4).

The distinction we draw between sites that currently meet the IUCN criteria (confirmed OECMs) versus those on-track to meet the criteria (candidate OECMs), highlights the existing ambiguity about where to draw the line between fully and partially addressing the relevant criteria (i.e., language used by the site-level assessment tool^12^). Our analysis found that most sites scored in the intermediate range for several of the indicators we developed to translate the broad criteria into a more granular assessment. This highlights how difficult it can be to determine whether sites fully or partially address the relevant criteria as currently described in the guidance. Because our indicators allow independent assessment of governance structures and aspects of management (Criterion 5), we were able to pinpoint important deficiencies in sites (Figures 6 & S3). A lack of resources and an inability to effectively manage threats to biodiversity (Criterion 6; Figure 5) are widespread challenges for area-based conservation^9,10,11,30^. While there has been considerable optimism that OECMs could be a mechanism to gain access to additional resources and build the capacity of sites for effective management^16^, there is currently no clear process to achieve this.

Demonstrating that sites currently meet the criteria for in-situ conservation of biodiversity requires evidence^12,31^. Providing the necessary evidence for effective biodiversity outcomes has been recognized as a potential barrier for sites seeking recognition as OECMs^13^. We found that a significant proportion of sites in our sample did not have adequate monitoring programs to demonstrate conservation outcomes (Figure S7). The site-level assessment tool encourages the use of documented evidence, but our data show that many sites will need to rely heavily on the knowledge of local experts. While considered a valid source of evidence for assessments^12^, the goal should be that monitoring data demonstrate a site is achieving positive conservation outcomes as an important safeguard to ensure OECMs deliver promised benefits^18^.

### Challenges for effective governance and management

Effective governance and management are strongly linked to positive conservation outcomes^9,11,30^. Threat management is clearly an issue for many sites, with most PAs and potential OECMs only partially addressing threats (Figure 6). In particularly, many sites were reported to be subject to activities that were at least somewhat incompatible with biodiversity conservation (Figure S4). Given that most potential OECMs had biodiversity conservation as a secondary objective (Figure 3), this may reflect a challenge in balancing management objectives so biodiversity outcomes are not compromised by other activities^32^. There has been a long-running debate in the PA literature about strictly protected versus multi-use areas that permit some resource extraction (e.g., ^32,33^). Ongoing monitoring will be critical to demonstrating sustainable use of natural resources over time.

Our results demonstrate that the capacity to achieve sustained conservation outcomes could be a major barrier to many sites meeting the OECM criteria. Many potential OECMs were considered unlikely to have the necessary structures in place to achieve long-term outcomes, often because they lack governance arrangements that offer long-term security (Figure 6). The legal frameworks that generally accompany PAs, while not infallible, are not easily overturned and offer a transparent mechanism through which changes to protection can be tracked (e.g., Protected Area Downgrading, Downsizing and Degazettement^34^). Most countries do not currently have specific legal mechanisms to recognize OECMs. Nor is there agreement about what types of governance instruments are required, or whether some existing legal structures are sufficient to be considered long-term (e.g.,^19^). Not only is the lack of formal structures a challenge for initially meeting the OECM criteria, but this also creates challenges for reporting and tracking the status of OECMs going forward.

The costs of demonstrating sites meet the requirements for OECMs could be prohibitive for Indigenous Peoples, local communities and other non-state actors^13^. Where sites do not currently meet the OECMs criteria, but could be on-track to do so, additional resources will be needed to help develop or improve the management and governance structures needed to assist sites meet their conservation goals. Our study suggests that many potential OECMs will be in this category, and there will need to be processes to support capacity building and develop incentive to ensure biodiversity outcomes can be maintained long-term. Without a dedicated source of funding for OECMs, there is a risk that expanding the area under conservation will increase existing capacity shortfalls^11^, and result in resources being diverted away from current PA management efforts^35^. Identifying innovative solutions to sustainably resource OECMs, and support potential OECMs to achieve the necessary standard, is therefore critical to ensuring the success of area-based conservation more generally.

### The role of OECMs in achieving ambitious area-based conservation targets

An important finding of our study is that potential OECMs were less likely than PAs to be of sufficient size to achieve the persistence of relevant species and ecosystems (Figure S6). Nevertheless, OECMs could still play an important role in buffering, enlarging or connecting existing PAs (Figure S2). Many PAs are too small to support sufficiently large populations to be viable long-term^36,37^, and have limited connectivity^38^, which may be a barrier to supporting biodiversity to adapt to changing climates. Therefore, considering OECMs in isolation may not reflect their true potential to contribute to the long-term conservation of biodiversity at ecologically-relevant scales. This emphasizes the importance of countries taking a holistic approach to area-based conservation as they move towards the 30% protection target in the Global Biodiversity Framework^6^. Supporting both OECMs and PAs to achieve effective conservation outcomes, as part of interconnected systems of conservation areas, will be an important part of ensuring meaningful benefits for biodiversity.

### Limitations and future directions

It is important to note that the sites included in our dataset, while reflecting diverse conservation areas around the world, were targeted because they meet the definition of potential OECMs - important places for biodiversity conservation. Therefore, the finding that a third of sites could be considered candidate OECMs may offer an overly optimistic assessment of the capacity for other, less well-supported conservation areas to meet the necessary criteria. There is likely to be a much wider spectrum of sites with different strengths and challenges. A more realistic approach could be to assess potential OECMs broadly, based on their capacity to meet the CBD criteria if given the right support (i.e., as candidate OECMs). This could be done using the more granular indicators we developed to identify where to target additional resources and technical advice to sites that have the potential to make important contributions to biodiversity conservation.

While local expert knowledge is an accepted means of verification for site-level assessments^12^, it is important to recognize that our evaluation did not collect quantitative data or consistently seek independent verification to support the assessments made by local experts. The indicators we developed are the most comprehensive attempt to transform the general CBD criteria into measurable attributes. An important next step must be determining what evidence is required to independently verify these assessments and demonstrate positive conservation outcomes, ideally with some quantitative metrics and clear thresholds for when a site has met a criterion.

While our study provides the most comprehensive evaluation of potential OECMs to date, our findings bring into focus the many questions that remain unanswered. It is incumbent on conservation science to develop the evidence-base required to understand what types of areas should be considered OECMs, what is required to demonstrate sites achieve the necessary criteria, and to support a wide range of sites to achieve and maintain conservation outcomes long-term. It is essential that the conservation community engage with these questions now. Our indicators provide an important tool to help make that leap. As we move towards 2030, we urge a renewed emphasis on ensuring area-based conservation is effective at protecting biodiversity. This will involve a deeper consideration of the role OECMs could play in supporting PAs to conserve important biodiversity as part of integrated conservation landscapes and seascapes.

## SUPPLEMENTAL INFORMATION

Appendix S1. Questionnaire,

Appendix S2 – Extended data - Figures S1–S9

## ACKNOWLEDGMENTS

We thank the many local experts who contributed their expertise to data collection: Gaspard Abitsi, Hugo Costa, Padu Franco, Saw Htun, Steve Insley, Nykol Jara, Devcharan (Dev) Jathanna, Khalil Karimov, Martin Mendez, Jean Mensa, Prayekti Ningtias, Joshua Pandong, Mauricio Palacios, Killivallavan Rayar, Martin Robards, Ravaka Ranaivoson, Ana Yi. The authors also acknowledge the contributions made by Indigenous peoples, local communities, local non-governmental organizations, government agencies and other local experts who have contributed their knowledge and understanding to support the site-level assessments. Special thanks to Jeanne Brown and Mel Berran for assistance with data collection and compilation.

## Notes

### Competing Interest Statement

The authors have declared no competing interest.

## REFERENCES

1. Watson, J.E.M., Dudley, N., Segan, D.B., and Hockings, M. (2014). The performance and potential of protected areas. Nature 515, 67–73. 10.1038/nature13947.

2. Maxwell, S.L., Cazalis, V., Dudley, N., Hoffmann, M., Rodrigues, A.S.L., Stolton, S., Visconti, P., Woodley, S., Kingston, N., Lewis, E., et al. (2020). Area-based conservation in the twenty-first century. Nature 586, 217–227. 10.1038/s41586-020-2773-z.

3. Convention on Biological Diversity (CBD) (2010). Strategic plan for biodiversity 2011–2020 and the Aichi targets. Report of the Tenth Meeting of the Conference of the Parties.

4. Intergovernmental Panel on Biodiversity and Ecosystem Services (IPBES) (2019). Summary for policymakers of the global assessment report on biodiversity and ecosystem services of the Intergovernmental Science-Policy Platform on Biodiversity and Ecosystem Services.

5. Dinerstein, E., Olson, D., Joshi, A., Vynne, C., Burgess, N.D., Wikramanayake, E., Hahn, N., Palminteri, S., Hedao, P., Noss, R., et al. (2017). An Ecoregion-Based Approach to Protecting Half the Terrestrial Realm. Bioscience 67, 534–545.

6. Convention on Biological Diversity (CBD) (2022). Kunming-Montreal Global Biodiversity Framework: COP 15 Decision L25.

7. Convention on Biological Diversity (CBD) (2018). Protected areas and other effective area-based conservation measures (Decision 14/8).

8. IUCN and World Commission on Protected Areas (IUCN-WCPA) Task Force on OECMs (2019). Recognising and Reporting Other Effective Area-based Conservation Measures. IUCN, Gland.

9. Geldmann, J., Barnes, M., Coad, L., Craigie, I.D., Hockings, M., and Burgess, N.D. (2013). Effectiveness of terrestrial protected areas in reducing habitat loss and population declines. Biol Conserv 161, 230–238. 10.1016/j.biocon.2013.02.018.

10. Edgar, G.J., Stuart-Smith, R.D., Willis, T.J., Kininmonth, S., Baker, S.C., Banks, S., Barrett, N.S., Becerro, M.A., Bernard, A.T.F., Berkhout, J., et al. (2014). Global conservation outcomes depend on marine protected areas with five key features. Nature 506, 216–220. 10.1038/nature13022.

11. Coad, L., Watson, J.E.M., Geldmann, J., Burgess, N.D., Leverington, F., Hockings, M., Knights, K., and Di Marco, M. (2019). Widespread shortfalls in protected area resourcing undermine efforts to conserve biodiversity. Front Ecol Environ 17, 259–264. 10.1002/fee.2042.

12. IUCN and World Commission on Protected Areas (IUCN-WCPA) (2023). Site-level tool for identifying other effective area-based conservation measures (OECMs). IUCN, Gland.

13. Alves-Pinto, H., Geldmann, J., Jonas, H., Maioli, V., Balmford, A., Latawiec, A.E., Crouzeilles, R., and Strassburg, B. (2021). Opportunities and challenges of other effective area-based conservation measures (OECMs) for biodiversity conservation. Perspect Ecol Conserv 19, 115–120. 10.1016/j.pecon.2021.01.004.

14. Protected Planet (2023). World Database on Other Effective area-based Conservation Measures. UNEP-WCMC. Accessed May 2023. https://protectedplanet.org.

15. Lemieux, C.J., Gray, P.A., Devillers, R., Wright, P.A., Dearden, P., Halpenny, E.A., Groulx, M., Beechey, T.J., and Beazley, K. (2019). How the race to achieve Aichi Target 11 could jeopardize the effective conservation of biodiversity in Canada and beyond. Mar Policy 99, 312–323. 10.1016/j.marpol.2018.10.029.

16. Gurney, G.G., Darling, E.S., Ahmadia, G.N., Agostini, V.N., Ban, N.C., Blythe, J., Claudet, J., Epstein, G., Estradivari, Himes-Cornell A., et al. (2021). Biodiversity needs every tool in the box: use OECMs Comment. Nature 595, 646–649.

17. Cook, C.N. (2023). Progress developing the concept of other effective area-based conservation measures. Conservation Biology. 10.1111/cobi.14106.

18. Dudley, N., Jonas, H., Nelson, F., Parrish, J., Pyhälä, A., Stolton, S., and Watson, J.E.M. (2018). The essential role of other effective area-based conservation measures in achieving big bold conservation targets. Glob Ecol Conserv 15, e00424. 10.1016/j.gecco.2018.e00424.

19. Marnewick, D., Stevens, C.M.D., Jonas, H., Antrobus-wuth, R., Wilson, N., and Theron, N. (2021). Assessing the extent and contribution of OECMs in South Africa. Parks 27, 57–70. 10.2305/IUCN.CH.2021.PARKS-27-1DM.en.

20. Dudley N, Robinson J, Andelman S, Bingham H, Conzo LA. (2022). Developing an outcomes-based approach to achieving Target 3 of the Global Biodiversity Framework. PARKS 28: 33–44.

21. Carroll, C., and Noss, R.F. (2022). How percentage-protected targets can support positive biodiversity outcomes. Conservation Biology 36, e13869. 10.1111/cobi.13869.

22. Maron, M., Simmonds, J.S., and Watson, J.E.M. (2018). Bold nature retention targets are essential for the global environment agenda. Nat Ecol Evol 2, 1194–1195. 10.1038/s41559-018-0595-2.

23. Jonas, H.D., MacKinnon, K., Dudley, N., Hockings, M., Jessen, S., Dan Laffoley, MacKinnon D., Matallana-Tobón, C.L., Sandwith, T., Waithaka, J., et al. (2018). Other effective area-based conservation measures: From Aichi Target 11 to the post-2020 biodiversity framework. Parks 24, 9–16. 10.2305/IUCN.CH.2018.PARKS-24-SIHDJ.en.

24. Pressey, R.L., Visconti, P., McKinnon, M.C., Gurney, G.G., Barnes, M.D., Glew, L., and Maron, M. (2021). The mismeasure of conservation. Trends Ecol Evol 36, 808–821. 10.1016/j.tree.2021.06.008.

25. Donald, P.F., Buchanan, G.M., Balmford, A., Bingham, H., Couturier, A.R., de la Rosa Jr., G.E., Gacheru, P., Herzog, S.K., Jathar, G., Kingston, N., et al. (2019). The prevalence, characteristics and effectiveness of Aichi Target 11’s “other effective area-based conservation measures” (OECMs) in Key Biodiversity Areas. Conserv Lett 12, e12659. 10.1111/conl.12659.

26. Jonas, H.D., Lee, E., Jonas, H.C., Matallana-Tobon, C., Wright, K.S., Nelson, F., and Enns, E. (2017). Will “Other Effective Area-based Conservation Measures” increase recognition and support for ICCAs? Parks 23, 63–78. 10.2305/iucn.ch.2017.parks-23-2hdj.en.

27. Mitchell, B.A., Fitzsimons, J.A., Stevens, C.M.D., and Wright, D.R. (2018). PPA or OECM? Differentiating between privately protected areas and other effective area-based conservation measures on private land. Parks 24, 49–60. 10.2305/IUCN.CH.2018.PARKS-24-SIBAM.en.

28. Rodríguez-Rodríguez, D., Sánchez-Espinosa, A., and Abdul Malak, D. (2021). Potential contribution of OECMs to international area-based conservation targets in a biodiversity rich country, Spain. J Nat Conserv 62. 10.1016/j.jnc.2021.126019.

29. Dudley, N. (2008). Guidelines for Applying Protected Area Management Categories. IUCN, Gland.

30. Gill, D.A., Mascia, M.B., Ahmadia, G.N., Glew, L., Lester, S.E., Barnes, M., Craigie, I., Darling, E.S., Free, C.M., Geldmann, J., et al. (2017). Capacity shortfalls hinder the performance of marine protected areas globally. Nature 543, 665–669. 10.1038/nature21708.

31. Geldmann, J., Deguignet, M., Balmford, A., Burgess, N.D., Dudley, N., Hockings, M., Kingston, N., Klimmek, H., Lewis, A.H., Rahbek, C., et al. (2021). Essential indicators for measuring site-based conservation effectiveness in the post-2020 global biodiversity framework. Conserv Lett 14, e12792 10.1111/conl.12792.

32. Leberger, R., Rosa, I.M.D., Guerra, C.A., Wolf, F., and Pereira, H.M. (2020). Global patterns of forest loss across IUCN categories of protected areas. Biol Conserv 241, 108299. 10.1016/j.biocon.2019.108299.

33. Hajjar, R., Oldekop, J.A., Cronkleton, P., Newton, P., Russell, A.J.M., & Zhou, W. (2021). A global analysis of the social and environmental outcomes of community forests. Nature Sustainability, 4, 216–224.

34. Golden Kroner, R.E., Qin, S., Cook, C.N., Krithivasan, R., Pack, S.M., Bonilla, O.D., Cort-Kansinally, K.A., Coutinho, B., Feng, M., Martínez Garcia, M.I., et al. (2019). The uncertain future of protected lands and waters. Science 364, 881–886.

35. Watson, J.E.M., Darling, E.S., Venter, O., Maron, M., Walston, J., Possingham, H.P., Dudley, N., Hockings, M., Barnes, M., and Brooks, T.M. (2016). Bolder science needed now for protected areas. Conservation Biology 30, 243–248. 10.1111/cobi.12645.

36. Clements, H.S., Kearney, S.G., and Cook, C.N. (2018). Moving from representation to persistence: The capacity of Australia’s National Reserve System to support viable populations of mammals. Divers Distrib 24, 1231–1241. 10.1111/ddi.12759.

37. Ivanova, I.M., and Cook, C.N. (2023). Public and private protected areas can work together to facilitate the long-term persistence of mammals. Environ Conserv 50, 58–66. 10.1017/s0376892922000455.

38. Ward, M., Saura, S., Williams, B., Ramírez-Delgado, J.P., Arafeh-Dalmau, N., Allan, J.R., Venter, O., Dubois, G., and Watson, J.E.M. (2020). Just ten percent of the global terrestrial protected area network is structurally connected via intact land. Nat Commun 11, 4563. 10.1038/s41467-020-18457-x.

39. Jones, K.R., Venter, O., Fuller, R.A., Allan, J.R., Maxwell, S.L., Negret, P.J., and Watson, J.E.M. (2018). One-third of global protected land is under intense human pressure. Science 360, 788–791.

40. Wildlife Conservation Society (WCS) (2023). 2023 Impact Report. WCS, New York. https://wcs.org/about-us/impact-report

41. Quinn, G.P., and Keough, M.J. (2002). Experimental design and data analysis for biologists (ProQuest).

