## Supplemental Information for "Evaluating the likelihood for areas important for conservation to be recognized as Other Effective area-based Conservation Measures"

### Appendix S1: Questionnaire

### Appendix 2 - Extended data

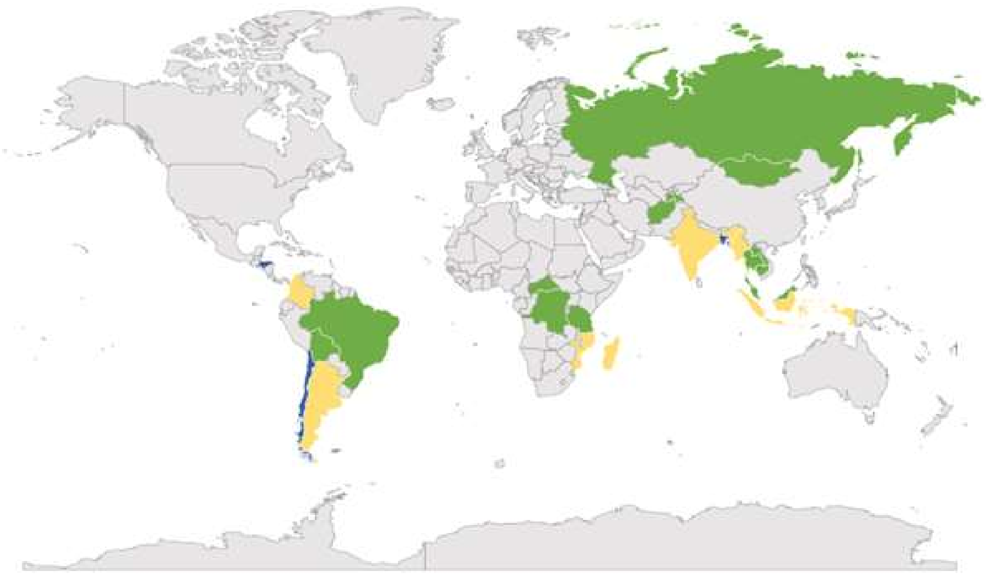

**Figure S1. Geographic distribution of sites across realms.**

Countries in blue are solely represented by marine sites (n=3). Countries in green are solely represented by terrestrial sites (n=13). Countries in yellow are represented by sites in both realms (n=7).

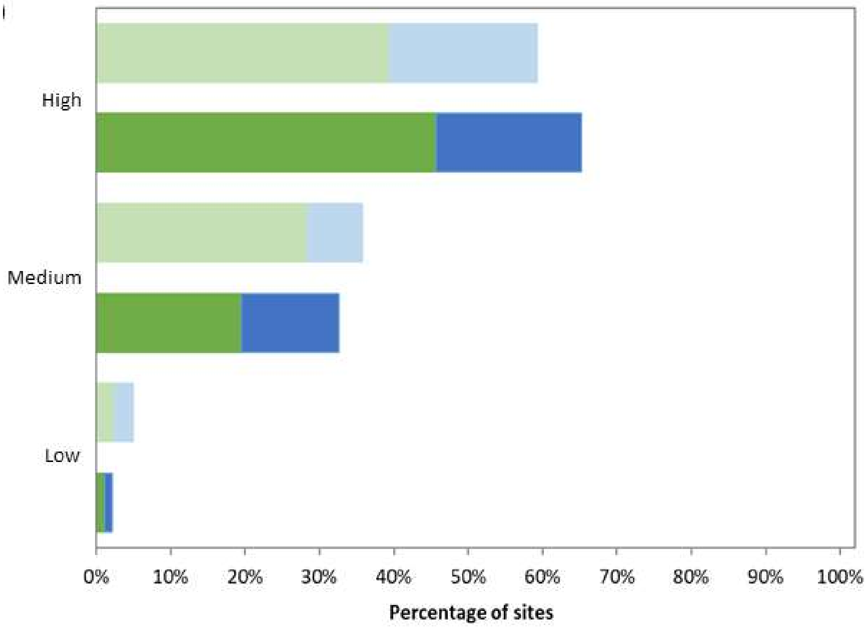


**Figure S2. The percentage of sites where ecological connectivity is high.**
Light bars = potential OECMs (n=81); Dark bars = PAs (n=92). Green = terrestrial sites (n=118); Blue = marine sites (n=55).

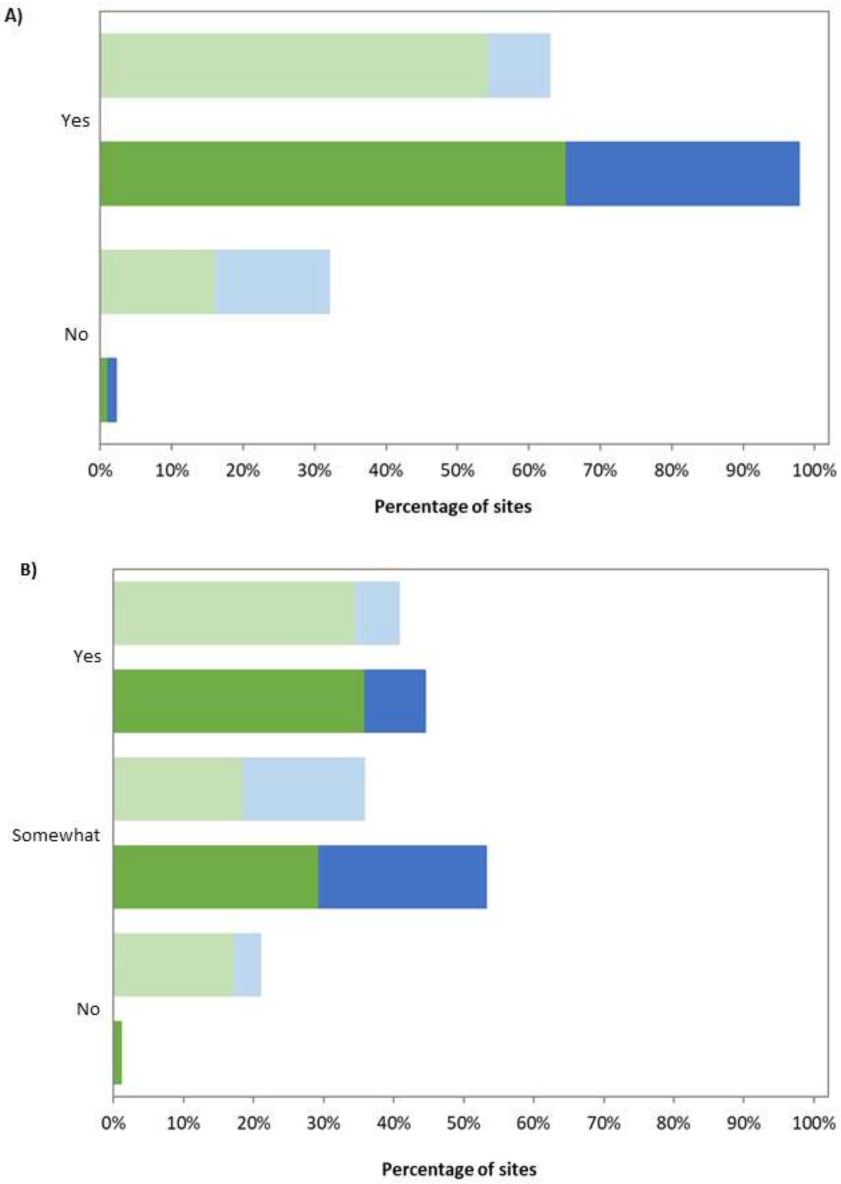


**Figure S3. The percentage of sites where governance authorities (A) have the necessary recognition to ensure that in-situ conservation of biodiversity can be achieved and (B) recognize and support its conservation-based status or the conservation-focused measures.**

Light bars = potential OECMs (n=81); Dark bars = PAs (n=92). Green = terrestrial sites (n=118); Blue = marine sites (n=55).

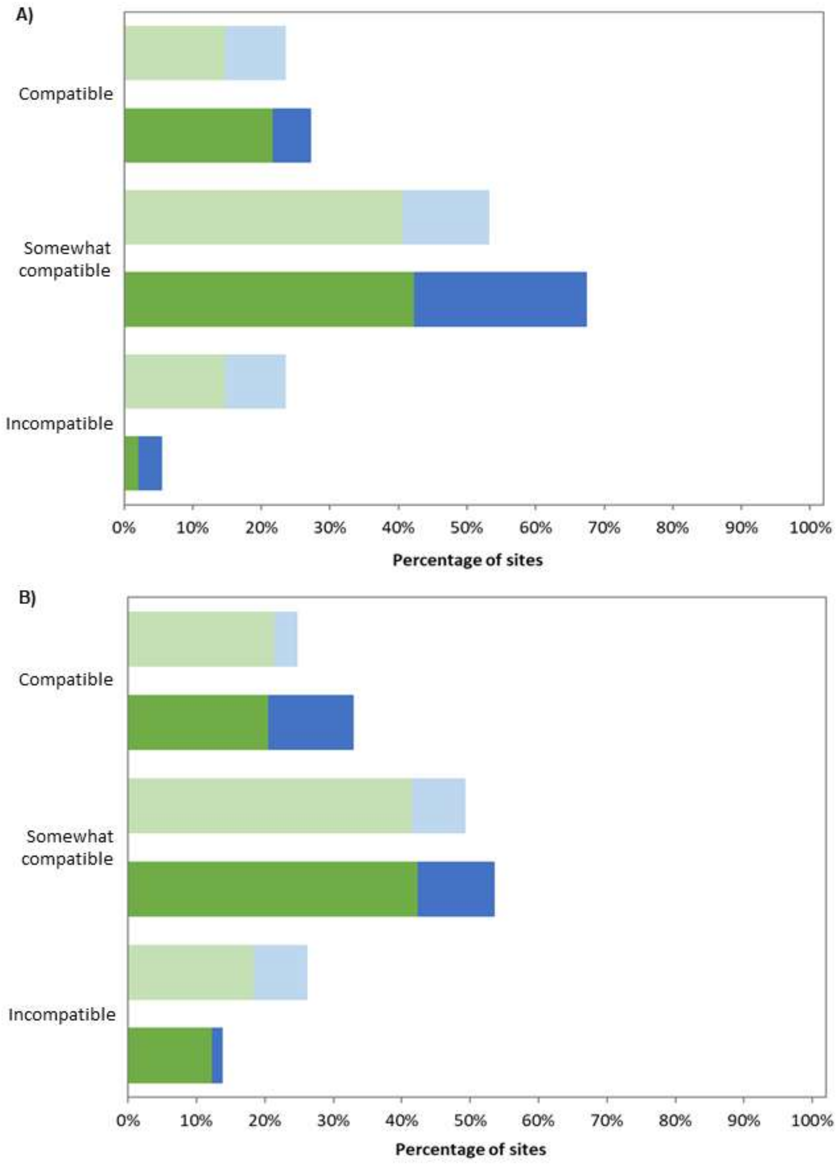


**Figure S4. The percentage of sites where activities are compatible with biodiversity conservation based on (A) subsistence and commercial uses, and (B) extractive industry and infrastructure.**

Light bars = potential OECMs (n=81); Dark bars = PAs (n=92). Green = terrestrial sites (n=118); Blue = marine sites (n=55).

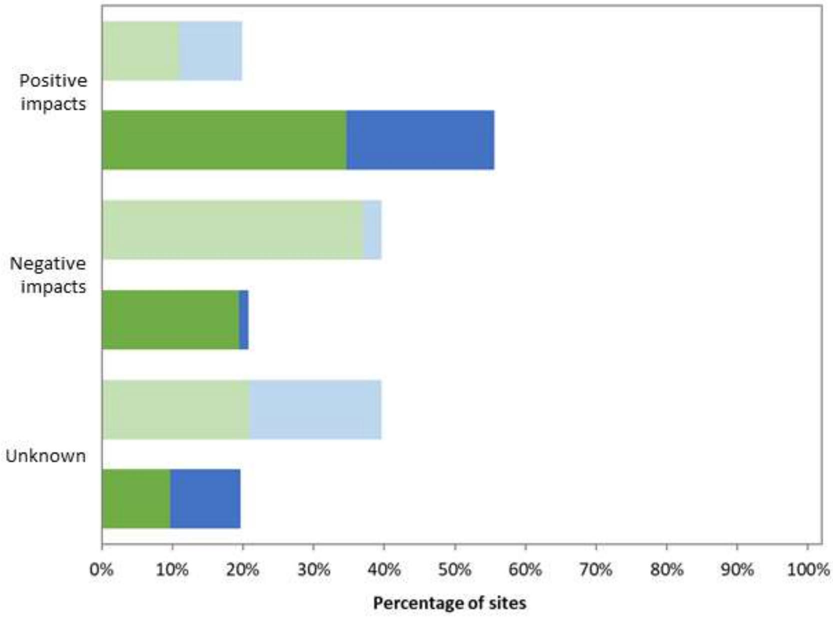


**Figure S5. The percentage of sites with government policies on natural resource management that support biodiversity.**

Light bars = potential OECMs (n=81); Dark bars = PAs (n=92). Green = terrestrial sites (n=118); Blue = marine sites (n=55)

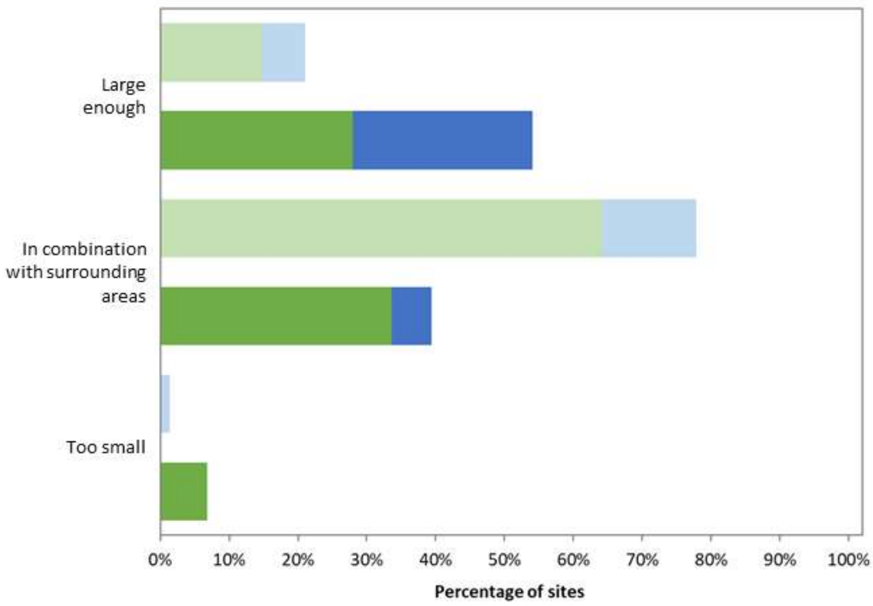


**Figure S6. The percentage of sites which are large enough to support biodiversity long-term.**

Light bars = potential OECMs (n=81); Dark bars = PAs (n=92). Green = terrestrial sites (n=118); Blue = marine sites (n=55).

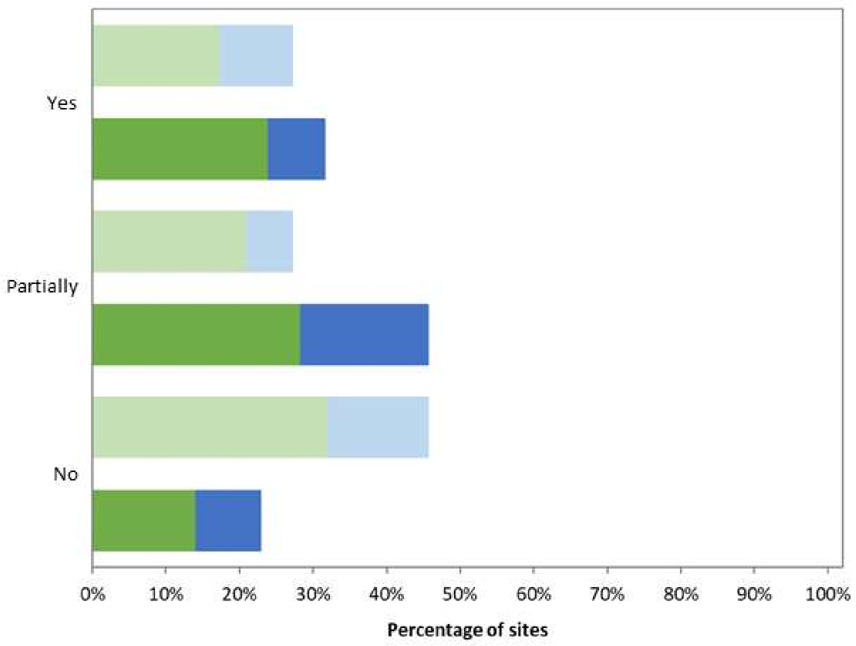


**Figure S7. The percentage of sites where monitoring of biodiversity outcomes is taking place.**

Light bars = potential OECMs (n=81); Dark bars = PAs (n=92). Green = terrestrial sites (n=118); Blue = marine sites (n=55).

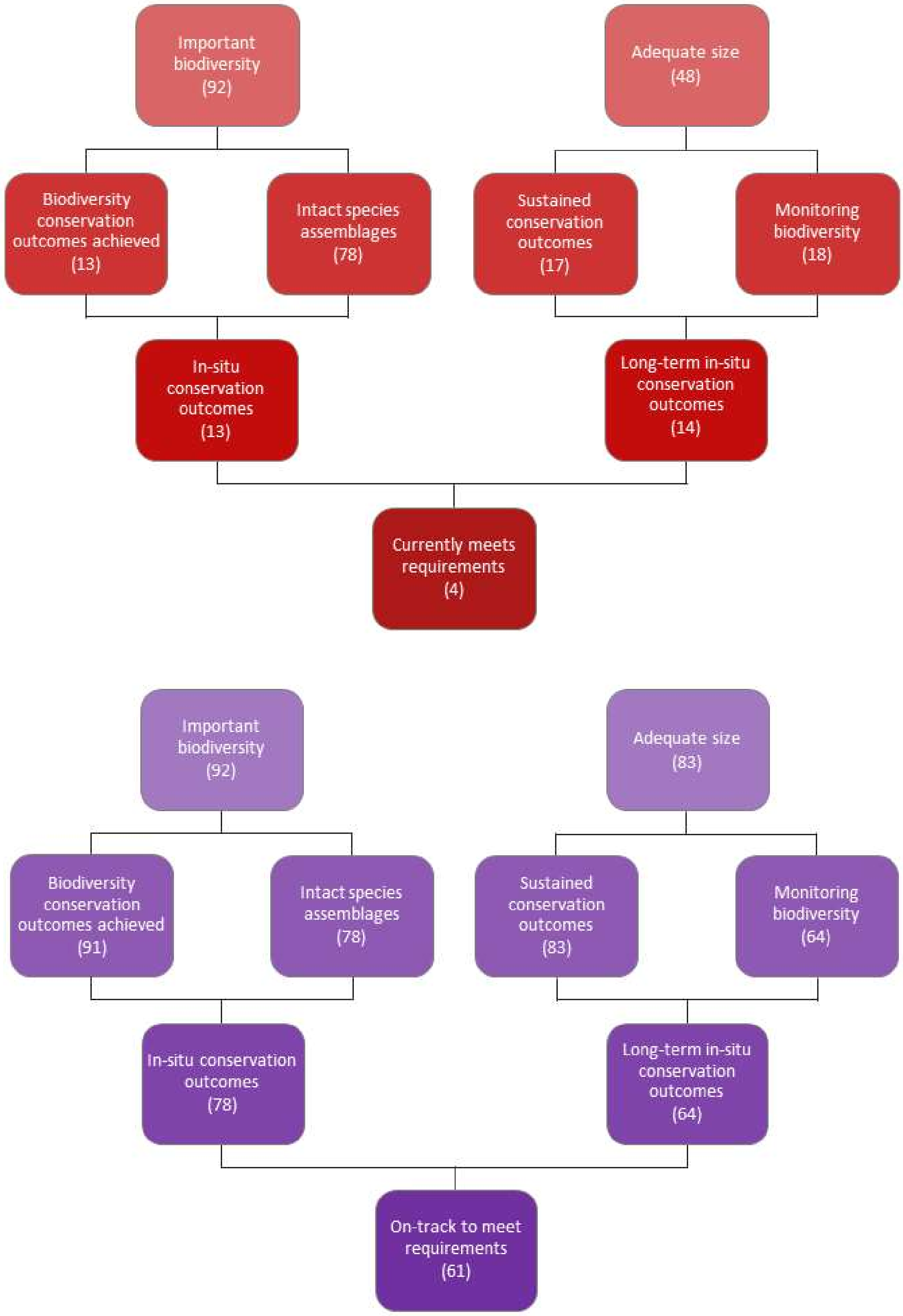


**Figure S8. The number of protected areas that currently (Panel A) or are on track to (Panels B) achieve effective and sustained in-situ conservation of biodiversity.**

Sites in Panel A could be equivalent to OECMs and sites in Panel B could be considered candidate OECMs. Numbers denote the number of sites, with 92 protected areas in total.

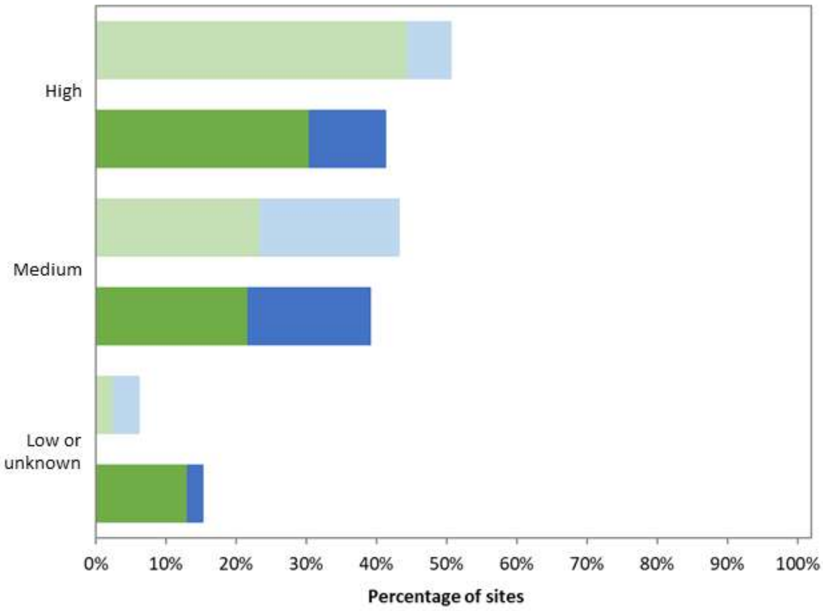


**Figure S9. The percentage of sites with high potential for restoration of biodiversity.**
Light bars = potential OECMs (n=81); Dark bars = PAs (n=92). Green = terrestrial sites (n=118); Blue = marine sites (n=55).
